# Recent evolution of a maternally-acting sex-determining supergene in a fly with single-sex broods

**DOI:** 10.1101/2022.11.24.517840

**Authors:** Robert B. Baird, John M. Urban, Andrew J. Mongue, Kamil S. Jaron, Christina N. Hodson, Malte Grewoldt, Simon H. Martin, Laura Ross

## Abstract

Sex determination is a key developmental process, yet it is remarkably variable across the tree of life. The dipteran family Sciaridae exhibits one of the most unusual sex determination systems in which mothers control offspring sex through selective elimination of paternal X chromosomes. Whereas in some members of the family females produce mixed-sex broods, others such as the dark-winged fungus gnat *Bradysia coprophila* are monogenic, with females producing single-sex broods. Female-producing females were previously found to be heterozygous for a large X-linked paracentric inversion (X’), which is maternally inherited and absent from male-producing females. Here we assembled and characterized the X’ sequence. As close sequence homology between the X and X’ made identification of the inversion challenging, we developed a k-mer-based approach to bin genomic reads before assembly. We confirmed that the inversion spans most of the X’ chromosome (approximately 55Mb) and encodes around 3500 genes. Analysis of the divergence between the inversion and the homologous region of the X revealed that it originated very recently (<0.5 mya). Surprisingly, we found that the X’ is more complex than previously thought and is likely to have undergone multiple rearrangements that have produced regions of varying ages, resembling a supergene composed of evolutionary strata. We found functional degradation of around 7.3% of genes within the region of recombination suppression, but no evidence of accumulation of repetitive elements. Our findings provide an indication that sex-linked inversions are driving turnover of the strange sex determination system in this family of flies.

## Introduction

Sex is an ancient feature shared by most eukaryotes, yet the sex determination systems regulating the development of males and females vary widely among animals (1) and can undergo rapid changes (2). Why such a fundamental developmental process as sex determination is variable remains an outstanding question (3). Insects include many examples of this diversity and are therefore an excellent model for understanding changes in sex determination systems. While most insects have genetic sex determination mechanisms with distinct sex chromosomes, different chromosomes act as the sex chromosomes in different species, and species differ in whether males (XY and X0 systems) or females (ZW and Z0 systems) are the heterogametic sex, and in divergence between the sex chromosome pair (1, 4). There are also examples of complete loss of sex chromosomes, where sex is linked to ploidy differences (e.g. haplodiploidy) or elimination or silencing of paternally-derived chromosomes in males. Another remarkable case, where sex is determined chromosomally but in a way that fundamentally differs from the standard XX/XY or ZZ/ZW system is that of monogenic sex determination. Here, sex is determined by the genotype of the mother instead of that of the zygote: mothers are genetically predetermined to produce either only male offspring or only female offspring. Monogenic sex determination has evolved in three clades of flies (Diptera): blowflies (Chrysomyinae, 5, 6), gall midges (Ceccidomyiidae, 7–9) and fungus gnats (Sciaridae, 10–12). Little is known about control of sex determination in blowflies (13). However, in the fungus gnat and gall midge species in which karyotypes have been characterized, monogeny appears to be associated with chromosomal inversions (14–16). None of these inversions has yet been characterized and little is known about the nature and the molecular evolution of these regions. Neither the evolutionary history of monogeny, nor how selection acts on sex determining regions that occur outside the context of conventional sex chromosomes, is thus currently understood.

Suppression of recombination through chromosomal inversions occurs in some sex chromosomes (17), and several situations can favor a lack of recombination (18–21). Prevailing theory posits that this process involves selection for suppressed recombination between the sex-determining locus, and sexually antagonistic alleles maintained polymorphically at partially sex-linked loci, potentially encompassing increasingly large portions of the sex chromosome in a stepwise process (22–24). However, several alternative hypotheses have recently been proposed, including a role for local adaptation (19), regulatory evolution (20) and the build-up of deleterious mutations (21). Such inversions may create regions that never or rarely recombine with their homologous X- and Z-linked regions. This creates sex-specific transmission and ensures that the affected regions are always heterozygous, unlike autosomal inversions. Such regions are likely to accumulate adaptive mutations specific to one sex or the other (19). If the region completely fails to recombine, it is liable to accumulate deleterious mutations and transposable elements (25, 26). As a result, sex chromosomes undergo functional degradation (27) and become a reservoir for repetitive sequences (28–30).

In the present study we investigated an X-linked inversion associated with monogeny in the fungus gnat *Bradysia (Sciara) coprophila*. This species has been studied extensively since the 1920s (31) and has a complex chromosome inheritance system (**Figure 1**). Like all members of Sciaridae, it reproduces through paternal genome elimination, where males fail to transmit paternally-derived chromosomes to their offspring as they undergo several rounds of maternally-controlled chromosome elimination targeting the paternal genome (10, 11). In all studied members of the Sciaridae, the somatic cells of males have an X0 karyotype, while those of females are XX. However, sex is determined by maternally-controlled X-elimination during early embryogenesis rather than X-inheritance. All zygotes begin with three X chromosomes, one inherited from the mother and two from the father -the result of aberrant spermatogenesis involving the nondisjunction of the maternal sister chromatids in the second meiotic division. During the 7th-9th embryonic cleavage divisions, either one or both paternal X chromosomes are eliminated from somatic cells, resulting in the zygotes developing into females (XX) or males (X0), respectively. The eliminated X chromosomes fail to divide at anaphase and are left behind on the metaphase plate (32). Germ cells in both sexes eliminate a single paternal X during a resting stage later in development.

**Figure 1.**
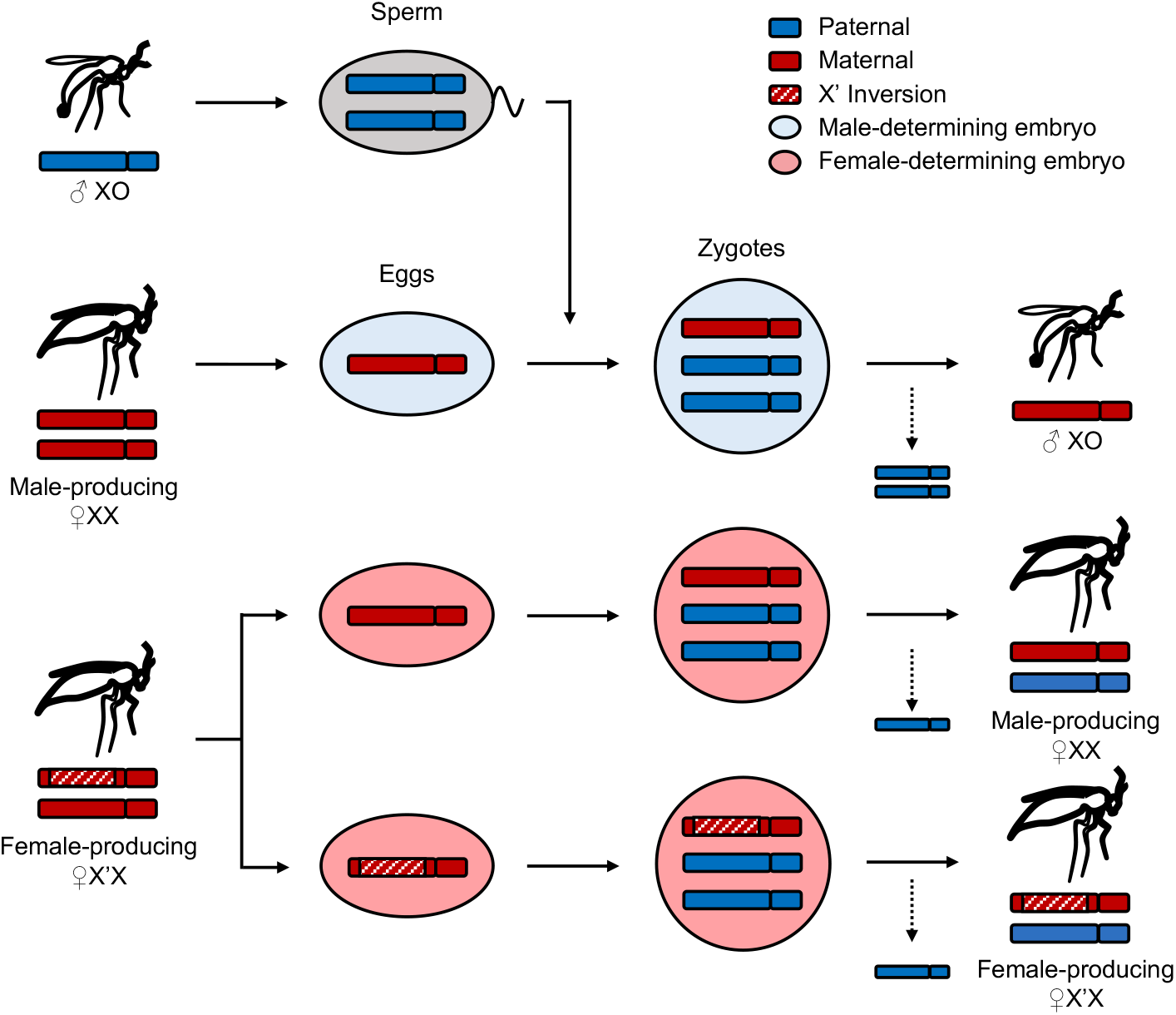
Sex determination and X chromosome inheritance in *B. coprophila*. While oogenesis is regular, sperm receive two X copies due to X nondisjunction. The mother’s genotype determines offspring sex: all zygotes begin with three X chromosomes and lose either one or two paternal X chromosomes via targeted paternal genome elimination, resulting in female and male development respectively. XX females produce only sons whereas females heterozygous for the X’ (X’X) produce only daughters.

In *B. coprophila* and many other Sciaridae, females are monogenic and produce single-sex progenies (33). Non-monogenic sciarids are ‘digenic’ and produce mixed-sex broods, although both monogenic and digenic species determine sex through X chromosome elimination (10). Both of these reproductive strategies occur in multiple Sciaridae genera (34, 35), though their evolutionary relationship to one another remains unclear. Early cytological observations suggested that two monogenic species, *B. coprophila* and *B. impatiens*, possess single long inversions spanning most of the X chromosome (henceforth the inverted chromosome is denoted by X’), and that female-producing females are heterozygotes (14, 36). Polytene chromosome staining indicates that such inversions are absent in digenic species (37, 38) as well as in at least one species exhibiting mixed reproductive strategies (39). Through a series of cytogenetic studies, Crouse (15, 36, 40, 41) deduced the structures of the chromosomes in *B. coprophila*, including the approximate breakpoint locations of the X’ inversion. Crouse (15) demonstrated that the inversion is paracentric and spans most of the length of X’, and that the two ends of the chromosome remain non-inverted and synapse with the X. The genome sequence of *B. coprophila*, with all three autosomes and the X chromosome, has recently become available (42, 43), though the sequence and nature of the X’ inversion remains unknown as the reference genome was generated from X0 males, which lack the X’ chromosome.

Here, we have shown through analysis of X-X’ divergence and variation that the structure of the X’ chromosome in *B. coprophila* is likely more complex than can be explained by a single paracentric inversion, and that it resembles a supergene composed of multiple linked inversions that all emerged less than 0.5 mya. Our finding that the X’ is young is intriguing given that monogenic reproduction is shared by multiple Sciaridae genera (10, 44), and suggests that inversions may drive the turnover of reproductive strategies in this family. We used a novel process of k-mer binning to assign short reads to chromosomes before assembly, allowing assembly of ∼55Mb corresponding to X’ supergene sequence despite its high sequence similarity to the ancestral X chromosome. With assembly and annotation of the X’ we compared patterns of evolution between the two homologous sequences which showed that the supergene shows some early signs of degradation characteristic of other evolving neo-sex chromosomes or neo-sex chromosome-like supergenes. We discuss the implications of our findings for disentangling the evolutionary relationship between the strange genetic properties of sciarid flies, and in light of the evolution of sex chromosomes and sex-linked adaptive inversions.

## Results

### X-X’ divergence reveals recent evolution and stratification of the X’ chromosome

We set out to identify the breakpoints of the long paracentric inversion previously described in the literature (36, 41). The size of the X chromosome in *B. coprophila* is estimated as 50 to 67 Mb (42, 45–47), and the X’ inversion spans almost the entire chromosome length (41). We therefore expected the inversion to be slightly shorter than the X. We produced whole-genome sequencing Illumina reads data from X0, XX, and X’X individuals, which when aligned against the recently updated chromosome-scale reference genome that contains sequences for chromosomes X, II, III and IV, but not X’ (Bcop_v2, 43), resulted in mapping rates of 93.58%, 96.68%, and 96.34%, respectively. That the X’X sample has approximately the same mapping rate as the XX sample, and higher than that for the X0 sample, indicates there is high enough sequence homology between the X and X’ to reliably call structural (SVs) and single-nucleotide variants (SNVs).

In an attempt to identify the breakpoints of the long paracentric inversion on the X’, we searched for SVs that could be attributed to the X’ using both Illumina short-read and PacBio long-read alignments from X’X samples, and using XX and X0 samples as a control. However, this analysis demonstrated that, in the X’X samples, the region of the X chromosome corresponding to the inverted region on the X’ is highly enriched for discordant paired-read and split long-read alignments that yield long, overlapping SV signals. We interpreted the entangled and contradictory nature of many individual SV calls as suggesting the presence of multiple complex rearrangements throughout the region rather than one single paracentric inversion (**Figure 2A, Table 1, Figure S1**). In contrast, HiC reads from X’X and X0 genotypes mapped against the X chromosome clearly revealed the two ‘main’ breakpoints observed cytologically (15), as well as three repeat regions that likely correspond to folds in the X chromosome (15, 41), but did not clearly show additional breakpoints along the chromosome (**Figure 2B,C**). Nonetheless, SNV calls revealed multiple distinct segments of the inversion with different SNV densities, again suggesting that multiple adjacent and/or nested inversions may have occurred at different times, perhaps in a stepwise fashion. We delineated putative evolutionary strata with a change-point analysis and used alignments of X’X Illumina reads against the X chromosome to estimate divergence for each stratum (**Figure 3, Table 2**). All strata were predicted to have emerged less than 0.5 mya. *Dxy* values calculated from all sites across the chromosome region were 0.0029 for the youngest stratum and 0.0071 for the oldest stratum, corresponding to divergence in years of 0.04 and 0.94 mya, respectively (assuming a similar mutation rate to *Drosophila*; see methods). Notably, some of the youngest strata had exceptionally low divergence. Estimates for neutrally evolving (synonymous) genic sites ranged from 0.07 to 0.34 mya for the youngest and oldest strata, respectively. Taken together, our findings suggest that a stepwise set of genomic rearrangements formed the X’ chromosome; we therefore set out to target the X’ sequence for *de novo* assembly.

**Table 1.** Number of each type of SV call from X0 and X’X alignments to the X chromosome.

| Support | SV | X0 genotype | X'X genotype |
| --- | --- | --- | --- |
| Short reads (Illumina) | Deletion | 20 | 2697 |
|  | Duplication | 1 | 56 |
|  | Inversion | 0 | 29 |
| Long reads (PacBio) | Deletion | 9 | 4037 |
|  | Duplication | 3 | 57 |
|  | Inversion | 11 | 267 |

**Table 2.** Divergence estimates for putative X’ strata in millions of years (mya). *Dxy* estimates are calculated from all sites along the portion of the X homologous to the supergene; neutral estimates are calculated from the proportion of synonymous variants in single-copy X-X’ homologs within each stratum. Lower (L) and upper (U) estimates are calculated from varying mutation rate and generation time estimates (see methods). Age estimates provided in the main text were calculated as mid-points between upper and lower estimates (**Figure 4B**).

| Stratum | Length (Mb) | *Dxy* | *Dxy* (mya, L) | *Dxy* (mya, U) | Neutral (mya, L) | Neutral (mya, U) | N homologs |
| --- | --- | --- | --- | --- | --- | --- | --- |
| S1 | 9.1 | 0.0050 | 0.033 | 0.097 | 0.075 | 0.219 | 326 |
| S2 | 0.4 | 0.0032 | 0.021 | 0.062 | 0.050 | 0.145 | 15 |
| S3 | 23.8 | 0.0066 | 0.044 | 0.128 | 0.113 | 0.329 | 904 |
| S4 | 1.0 | 0.0029 | 0.020 | 0.057 | 0.034 | 0.099 | 35 |
| S5 | 1.5 | 0.0068 | 0.045 | 0.133 | 0.145 | 0.424 | 50 |
| S6 | 0.1 | 0.0061 | 0.041 | 0.120 | NA | NA | 0 |
| S7 | 9.3 | 0.0070 | 0.047 | 0.137 | 0.150 | 0.436 | 351 |
| S8 | 0.1 | 0.0073 | 0.049 | 0.143 | 0.050 | 0.147 | 3 |
| S9 | 2.8 | 0.0067 | 0.045 | 0.131 | 0.160 | 0.467 | 65 |
| S10 | 0.4 | 0.0062 | 0.042 | 0.122 | 0.119 | 0.346 | 22 |
| S11 | 1.5 | 0.0069 | 0.046 | 0.135 | 0.141 | 0.412 | 50 |
| S12 | 1.9 | 0.0071 | 0.048 | 0.139 | 0.171 | 0.499 | 96 |
| S13 | 1.3 | 0.0069 | 0.046 | 0.134 | 0.132 | 0.386 | 90 |
| S14 | 1.2 | 0.0059 | 0.040 | 0.115 | 0.146 | 0.425 | 36 |
| S15 | 3.5 | 0.0056 | 0.038 | 0.109 | 0.075 | 0.218 | 97 |
| S16 | 0.9 | 0.0057 | 0.038 | 0.111 | 0.102 | 0.297 | 54 |
| **Total** | **58.8** | **0.0054** | **0.036** | **0.106** | **0.116** | **0.271** | **2204** |

**Figure 2.**
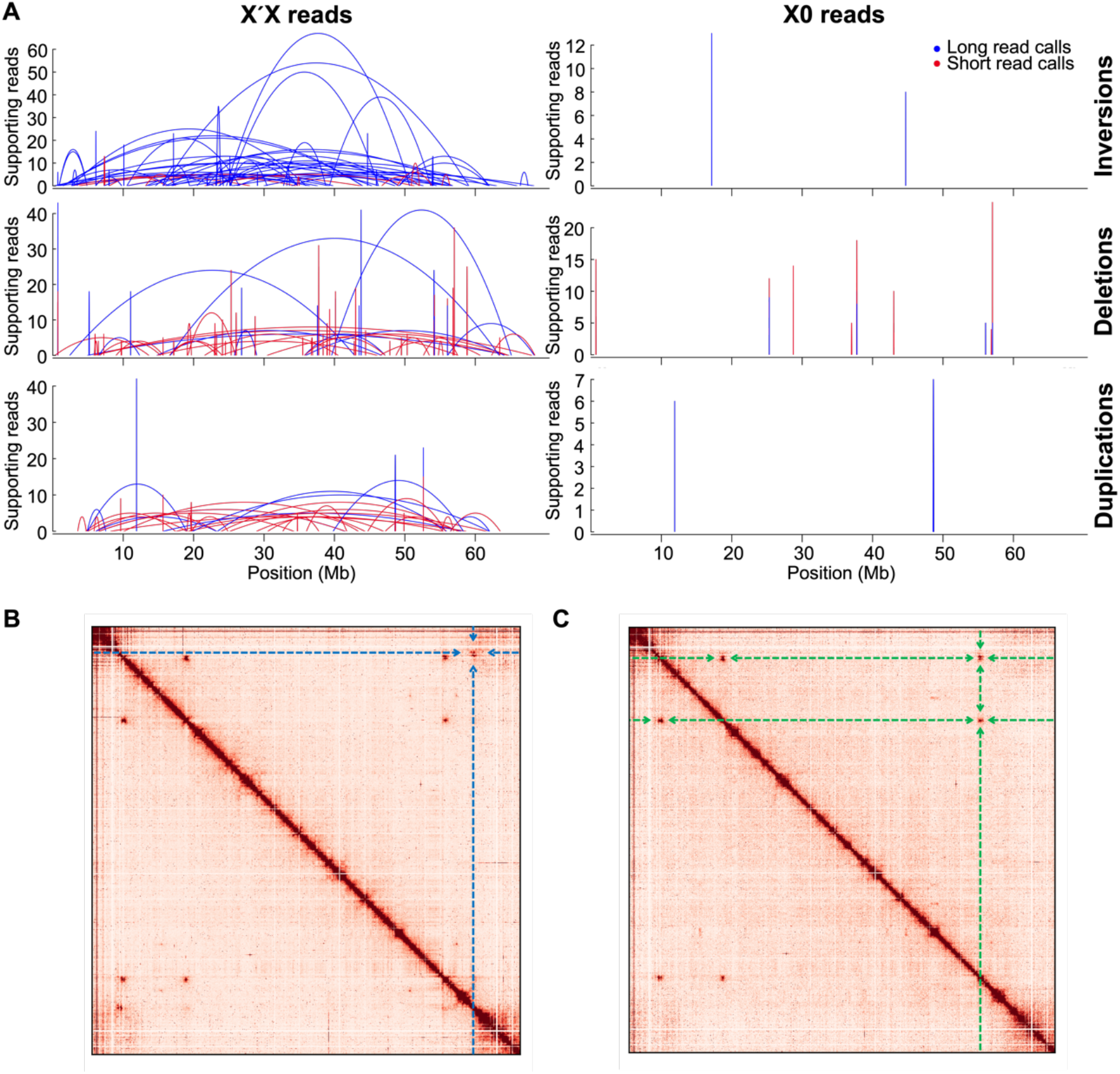
(A) SV calls from the X’X genotype are enriched across the X chromosome compared to calls from the X0 genotype, indicating more complex rearrangements for the X’ than may be explained by a single paracentric inversion. Start and end positions of SVs are shown with arcs. Only SVs supported by at least 4 reads and with spans greater than 10kb are shown. (B) HiC contact heatmap across the X chromosome for reads from the X’X genotype, as well as (C) for the X0 genotype. Contact showing the two main breakpoints is highlighted by blue dashed lines in (B). Repeats present in both heatmaps are highlighted by green dashed lines in (C).

**Figure 3.**
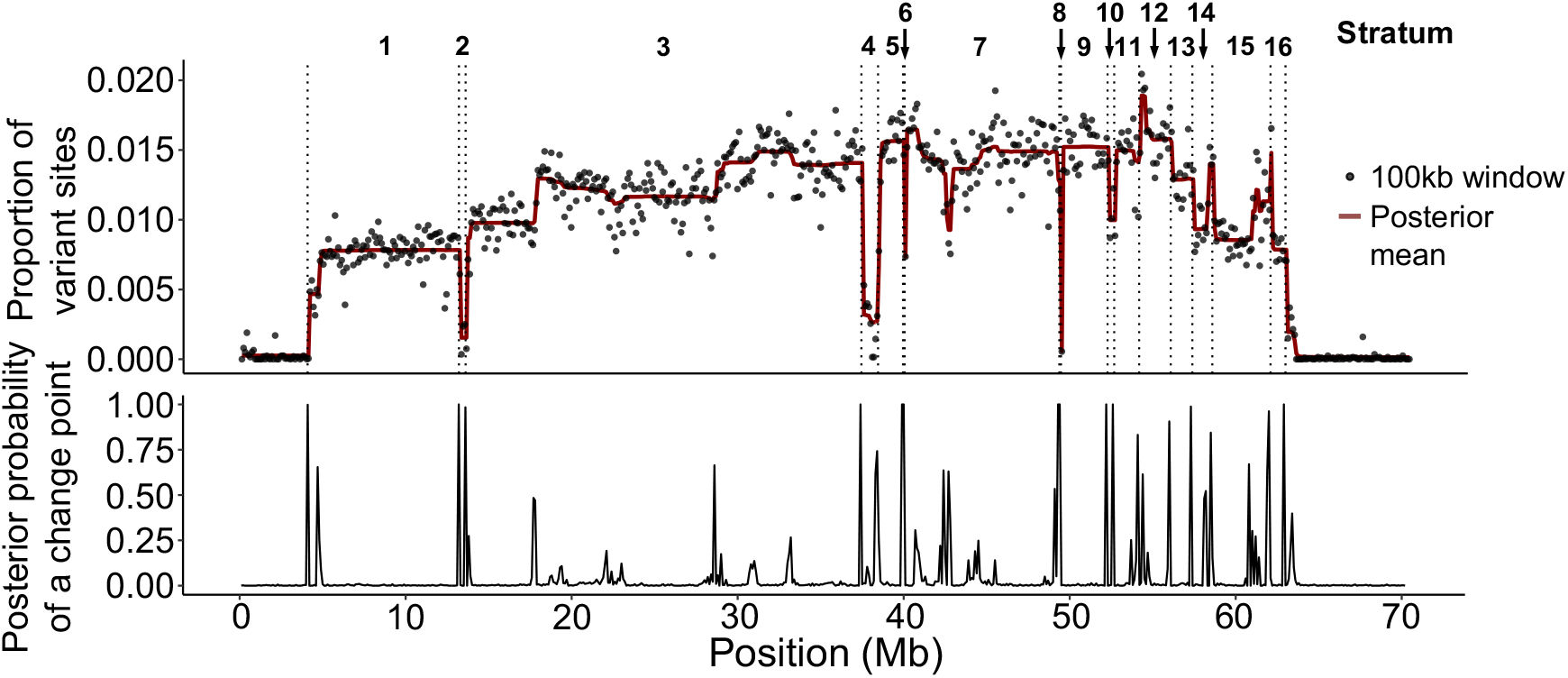
Upper panel: the distribution of variant sites between the X and X’, along which posterior means were calculated. Lower panel: the probability of point changes between posterior means were used to delineate putative evolutionary strata. Putative breakpoints between strata are shown as dotted lines in the upper panel.

### *De novo* assembly of the X’ sequence

We attempted assembly of PacBio reads from X’X individuals, followed by chromosome assignment of scaffolds using sex differences in read depth across the genome (48, **S1 Text, Figure S2**). This yielded a genome size of only 291Mb which was comparable to the size of the male (X0) genome (42). Moreover, we were able to assign only around 3.6Mb as putative X’ sequence (**Table S1**). High sequence homology between reads originating from the X and X’ chromosomes was likely leading to their collapsing together upon assembly. To overcome this we used a process akin to haplotype resolution of diploid sequences by trio binning (49). Our approach utilizes differences in k-mer frequencies in Illumina reads between sexes to assign them to chromosomes prior to assembly. Taking advantage of high homozygosity due to over a century of inbreeding (50) and the fact that X’ is limited to X’X individuals, we assigned k-mers specific to X’X female reads as likely to belong to the X’ (**Figure 4A**). We used these k-mers to extract the short reads from the X’X dataset as putative X’-specific reads. In contrast to long reads, which have a high likelihood of false k-mer matches due to high sequencing error rates, we found that short reads (75-150bp) can effectively be binned with k-mers due to their low error rate and short length (**S2 Text**).

**Figure 4.**
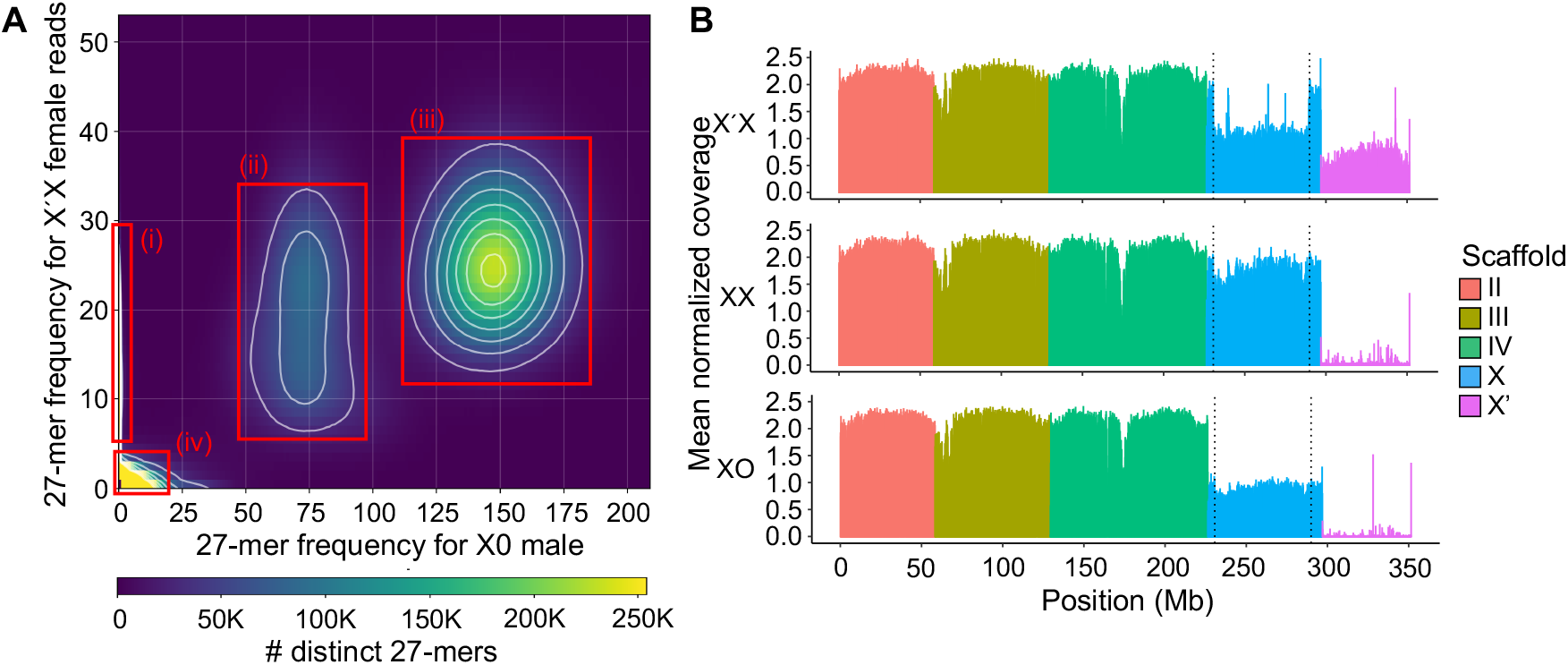
(A) 27-mer frequency heatmap between Illumina reads from X0 versus X’X flies. 27-mers form dense clouds based on k-mer frequency, which reflects ploidy: (i) 27-mers specific to X’X are assigned as putative X’-specific 27-mers. (ii) 27-mers haploid in X0 but diploid in X’X are likely those belonging to the X and the portion of X’ shared by both chromosomes. (iii) 27-mers diploid in both sexes are likely those belonging to autosomes. (iv) 27-mers containing read errors cluster around the origin. (B) Mean normalized per-based genomic coverage across 100Kb windows of all autosomes, the X and the X’-supergene sequence, for reads from pooled individuals of each genotype: X’X (top), XX (middle) and X0 (bottom). The main breakpoints of the X’ (dotted lines) are clearly visible in coverage from X’X reads.

The resulting putatively X’-specific Illumina reads assembled into 61.7Mb across 42564 contigs. We performed reference-based scaffolding using the X chromosome (43) to constitute a single scaffold corresponding to the X’ supergene. To gap-fill and polish the X’ scaffold, we combined it with the remaining chromosomes II, III, IV and X (43), then used PacBio reads from X’X individuals competitively mapped against all chromosomes to fill some remaining gaps (**Figure S3**, and Illumina reads from X’X individuals to polish the final assembly. The resulting 55.4Mb scaffold is the first model of the X’ sequence contained within the long paracentric inversion breakpoints defined by Crouse (15, **Table 3**). Due to using the uninverted X as a reference for scaffolding the X’ contigs, this scaffold may be an inaccurate structural representation of the X’ chromosome. However, alignment of Illumina reads from all three genotypes (X0, XX and X’X) to all chromosomes strongly supports its correspondence to the X’ chromosome (**Figure 4B**). As expected, we observed that the X’ sequence (i) had haploid coverage (1n) across the X’ sequence and the corresponding inverted region of the X chromosome compared to the autosomes for X’X individuals, (ii) had comparatively low coverage of the X’ for XX and X0 individuals, and (iii) had relatively equal coverage across the X and autosomes in XX females (**Figure 4B**).

**Table 3.** Assembly statistics for the *B. coprophila* chromosome-level genome (43), also showing the X’-supergene sequence. Predicted gene shown are those annotated across the entire genome in this study.

| Chromosome | Size (Mb, excluding Ns) | N count | Gaps | Predicted genes |
| --- | --- | --- | --- | --- |
| II | 57.99 | 57279 | 26 | 3968 |
| III | 69.01 | 2035696 | 101 | 5919 |
| IV | 95.49 | 1593141 | 90 | 6730 |
| X | 67.23 | 3282753 | 187 | 3881 |
| X'-supergene | 55.42 | 323958 | 2790 | 3470 |
| Total (excluding X') | 289.72 | 7289104 | 404 | 20498 |
| Total | 345.13 | 7613062 | 3194 | 23968 |

### Functional degradation of the X’ chromosome

We identified 3470 protein-coding genes totaling 4.2Mb, i.e. 7.6% of the 55.4 Mb X’ supergene scaffold. The portion of the X chromosome homologous to the supergene spans 58.8Mb and contains 3429 genes totaling 5.2Mb, i.e. 8.8% of the region. Thus, the proportion of the chromosome corresponding to coding sequence is slightly lower on the X’ supergene relative to the homologous X region, which may reflect an increase in non-coding DNA within the supergene. We identified 2321 single-copy homologs in both the regions. A further 527 genes from the X’ and 679 genes from the X across 296 orthologous groups (OGs) were categorized as duplicates. In 64 duplicate OGs, the X’ and X chromosomes had the same number of gene copies. Of the remaining duplicate OGs, 162 had more copies on the X compared to the X’, while 70 had more copies on the X’. These may be due to X- and X’-specific duplication events, or ancestral duplication and subsequent loss in one or the other chromosome. Unlike X mutations, X’ mutations should not be purged by recombination, thus deletions and duplications on the X’ are the more likely explanation. We also found that 622 genes across 603 OGs were specific to the X’, and 429 (across 359 OGs) were specific to the X. Gain of novel genes and whole-gene deletions from the X’ are possible explanations for these finding, although sequence divergence, misassemblies or gaps may have also led to homologs not being found.

We investigated the possibility that the X’ has undergone patterns of degeneration similar to other non-recombining sex chromosomes by analyzing functional degradation of genes and accumulation of repetitive elements. Of the 2321 single-copy gene OGs, we found that 123 (5.3%) contain X’-specific mutations that are likely to compromise gene function (including frameshift and/or gain or loss of stop or start codon mutations, **Table S2**). We further analyzed expression of single-copy homologs and found that fewer X’ genes are transcribed compared to their X-linked homologs: across four life stages in females, 2191 X’ homologs are expressed compared to 2237 X homologs, although only the larval stage had significantly fewer X’ copies expressed (**Figure 5A**). Overall, 7.3% of genes were classified as either silenced, disrupted or both (**Figure 5B**).

**Figure 5.**
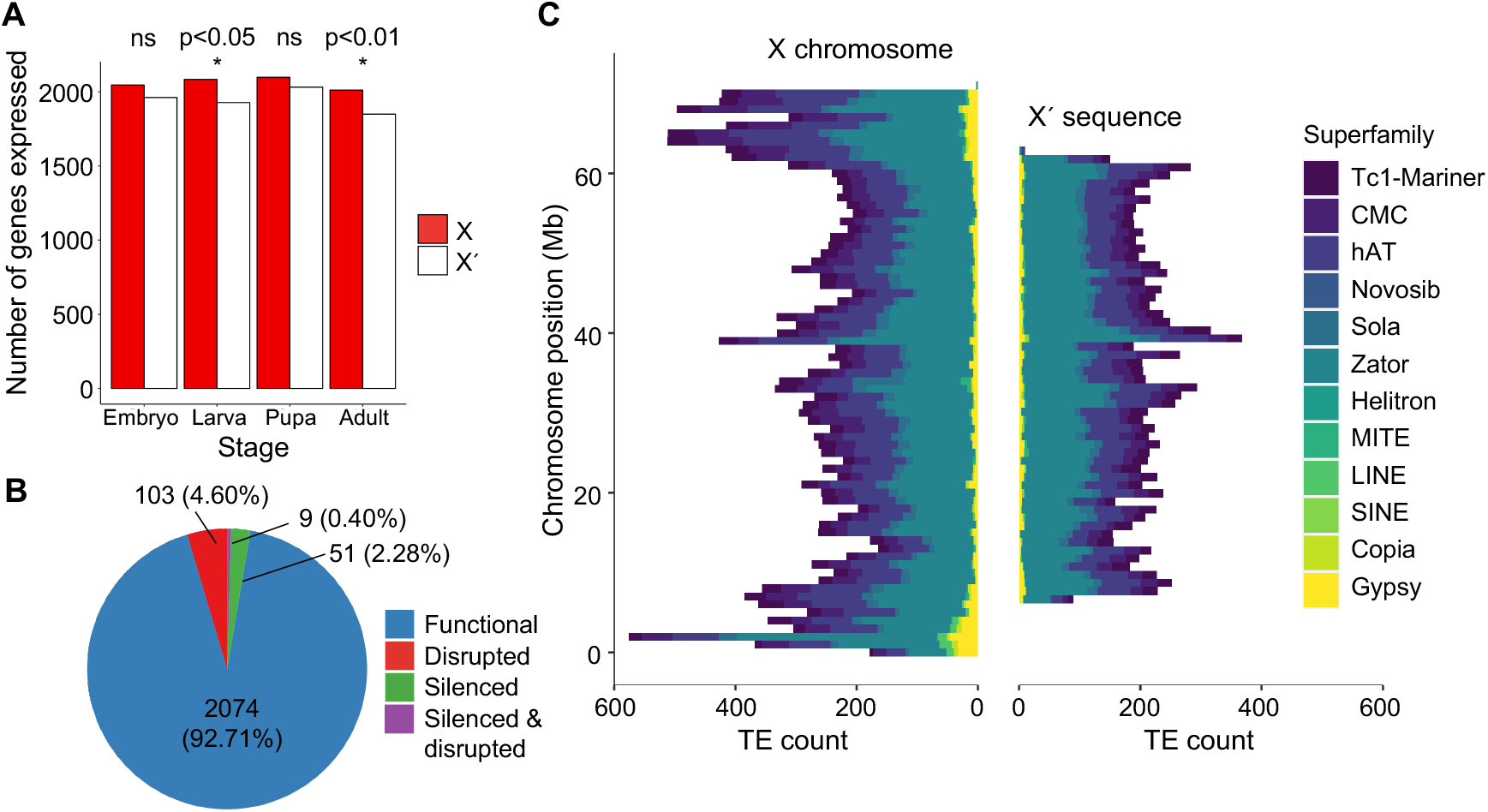
(A) Number of single-copy orthologous genes expressed at four developmental stages in females. Significantly more genes expressed from the X compared to the X’ chromosome in larvae (X-squared = 6.07, df = 1, p-value = 0.014) and adults (X-squared = 6.80, df = 1, p-value = 0.009); other stages were non-significant (ns). (B) Proportion of functional versus nonfunctional X’-linked genes that have functional X homologs. (C) TE counts in 1Mb bins distributed on the X chromosome and within the X’ supergene sequence.

The fly stock used in this study was derived from a laboratory stock maintained by Metz (51), in which X’X females carry an X’-linked, irradiation-induced mutation that causes *Wavy* wings. Among the genes classified as degraded, we identified several with functions in wing development, including the wing polarity protein STAN and the “held out wings” protein HOW, which are both required for regular wing development in *Drosophila* (52, 53). Since sex determination occurs via elimination of one, rather than two, X chromosomes in the embryos of X’X females, we may also expect that among those genes disrupted or silenced are sex-determination candidates. Among the genes classified as degraded, we found several candidates involved in chromatin regulation, chromosome segregation and cohesion (**S3 Text, Table S3**).

We also checked X’X females for evidence of dosage compensation (DC) of X-linked genes that have corresponding degraded X’ homologs. We expected that if these genes were dosage compensated, they would be upregulated in X’X females to match the expression of those genes in XX females where both copies are functional. We compared expression of X-linked genes in X’X and XX female samples (**Figure S4**. In total, only 4 X-linked genes were significantly upregulated in the X’X females (**Table S4**), none of which were genes that we classified as pseuogenized. Thus, we found no evidence of DC of degraded genes.

We also analyzed transposable element (TE) content across the two different X chromosomes and did not find an enrichment of repetitive sequences on the X’. In total, the TE count within the X’-specific sequence was 11982 compared to 16050 for the homologous portion of the X, corresponding to densities of 17.4% and 10.4% of the assembled sequence for each chromosome region, respectively. Moreover, the distribution of TEs and TE superfamilies was similar between the two chromosome regions (**Figure 5C**).

## Discussion

Recently-evolved sex chromosomes can provide crucial insights into understanding the evolution and turnover of sex chromosomes. *B. coprophila* is particularly unusual in that it exhibits a major transition from an X0-like system to one not unlike a ZW system, though only half of females are heterogametic and the sex chromosome is maternally-acting rather than acting in the zygote. Most known cases of neo-sex chromosome evolution in Diptera involve XY-XY transitions resulting from sex chromosome-autosome fusions (54), thus *Bradysia* offers a chance to study the early differentiation of the sex chromosomes in a unique evolutionary context. Our analysis and assembly of the ∼55Mb X’ sequence revealed that it evolved very recently (<0.5 mya), that it is more complex in structure than previously thought, and that it is beginning to undergo functional degradation characteristic of classical sex chromosome evolution.

### The X’ chromosome evolved recently and shows signs of degradation

In insects, sex chromosome turnovers and emergences of neo-sex chromosomes are common (55, 56), among which *Drosophila* neo-Y chromosomes are among the most well-studied (54, 57). In *Drosophila* males are achiasmatic, so Y-fused autosomes instantly become completely sex-linked and non-recombining (58). In the absence of recombination, deleterious mutations and TEs accumulate irreversibly because offspring will inherit the full mutational load of their parents (59). Furthermore, lack of recombination leads to hitchhiking effects of deleterious mutations, where beneficial mutations arising on the sex-limited chromosome spread to fixation and carry with them linked mildly deleterious loci (23, 60). The inevitable result of such processes is functional degradation of the non-recombining chromosomes (61, 62). Studies of neo-sex chromosome evolution in *Drosophila* have indeed found that significant degeneration occurs rapidly and correlates with age. In *D. pseudoobscura*, the neo-Y evolved ∼15 mya and has very few genes remaining (63, 64). The *D. miranda* neo-Y has undergone ∼4-fold expansion but 40% of the ancestral genes are pseudogenized via TE insertions, frameshifts or other nonsynonymous mutations in only ∼1.5 million years of evolution (27, 65–68). The younger *D. busckii* and *D. americana* neo-Ys evolved ∼0.8 and <0.47 mya and have 60% and 22% pseudogenized genes, respectively (57, 69, 70). The *D. albomicans* neo-Y chromosome, which is composed of several haplotype blocks that underwent recombination suppression separately between 0.09 and 0.14 mya, exhibits very few nonsynonymous changes (71). Our finding that around 7.3% of genes on the <0.5 mya *B. coprophila* X’ chromosome are pseudogenized is consistent with this expected trajectory of sex chromosome evolution. Moreover, because the chromosome is present in only half of females, its effective population size is half that of other sex-limited chromosomes. As such, it should exhibit an accelerated rate of decay as the effects of drift should increase the rate of evolution further at sites under purifying selection (72). To explore the potential effects of drift on sex-limited chromosome degeneration further, it will be essential to compare the X’ chromosomes of other Sciaridae species which may have evolved independently and at different times, such as that of *B. impatiens* (see below, 14).

Sex chromosome degeneration is sometimes accompanied by the evolution of dosage compensation (DC) mechanisms to re-establish diploid expression of the X chromosome in males (73). In the ancestral *Drosophila* X chromosome this is achieved by hypertranscription of X-linked genes in males (74). The neo-X chromosomes of *D. melanica* and *D. robusta*, which evolved independently 4-15 mya (75), achieved global DC via repeated transposon-mediated co-option of the motif required for recruitment of the DC complex that originally evolved to compensate the ancestral X (76). The younger *D. miranda* neo-X chromosome is in a transient state between gene-by-gene and global DC (77–79), and there is little evidence of any form of DC in the even younger *D. americana* and *D. albomicans* neo-X chromosomes (57). In *B. coprophila* there is evidence for DC in X0 males through upregulation of X expression (42, 80). As for the X’X females, we found no evidence that X-linked genes with degraded X’ homologs show DC in X’X females. The young age of the X’ may mean that there has not been sufficient time for the establishment of DC mechanisms to compensate for degraded X’ genes. Furthermore, because the X’ chromosome is present in only half of females, conflict between XX and X’X females over gene expression may hinder the evolution of DC.

Accumulation of repetitive sequences occurs rapidly in recently-evolved sex chromosomes (29, 81, 82). The neo-Y chromosome of *D. miranda* has undergone massive TE accumulation and has expanded significantly as a result (27, 68), though the repetitive landscape of younger (<0.5 mya) *Drosophila* neo-Ys (57) has not been analyzed and the speed at which TEs accumulate in non-recombining regions is unclear. Despite the recent divergence between X and X’ in *B. coprophila*, one would expect at least slightly higher TE content within the non-recombining region. However, TE abundances vary significantly even between closely related species: among *Drosophila* species the proportion of the genome composed of TEs varies between 10 and 40% (83). In birds, extensive TE accumulation is found on W chromosomes, despite low genome-wide TE content (84). Different lineages also have differing proportions of DNA transposons relative to retrotransposons. We find that *B. coprophila*, similar to the *Musca, Aedes* and *Culex* genera, has a higher proportion of DNA transposons than *Drosophila* (83). Compared to the cut-and-paste mechanisms of DNA transposons, the copy-and-paste mechanisms of retrotransposons may lend themselves to more rapid accumulation (85). Alternatively, the lack of TE accumulation on the X’ chromosome could be explained simply by the young age of the X’. However, it should be noted that our approach using X’-specific k-mers to assemble the X’ may result in failure to assemble repetitive sequences that are shared by other chromosomes, which may explain why we found a lower TE content on the X’ compared to the homologous X region. Understanding the dynamics of TE accumulation in this peculiar system will require a more contiguous assembly of the X’ as well as examination of other X’ chromosomes in Sciaridae species (see below).

### A role for adaptive stepwise expansion of the X’

Over the last decade, diverse and complex phenotypes that segregate within populations have been shown to be controlled by clusters of linked loci in inversions and/or inversion-based supergenes, as was proposed long ago for Batesian mimetic morphs of butterflies (86–88). Similar genetic systems have been discovered in other organisms, including adaptively evolving sex-linked genes in a moth (89, 90), divergent social behaviors in ants (91, 92), mating compatibility in fungi (93), and several polymorphisms in birds including plumage color (94), reproductive strategies (95) and sperm morphology (96). This study of the likely recent transition in the sex-determining system in flies presents another case. It has been argued that supergenes may be more widespread than previously recognised, that they are important for co-segregation of adaptive variation within a species, and that they may even occasionally result in the spread of complex phenotypes across species boundaries (97, 98).

The evolutionary trajectories of supergenes and sex chromosomes show similarities: some supergenes have evolved in a stepwise manner or undergone functional degradation, and sex chromosomes also play an important role in adaptation and speciation (93, 99–102). Furthermore, the evolutionary fates of inversions differ depending on whether they arise on sex chromosomes or on autosomes, with the probability of spread of an inversion through a population being higher on sex chromosomes. This is further affected by sex-biased migration patterns, dominance of locally adapted alleles and chromosome-specific deleterious mutation load (19). Indeed, X-linked genes are predicted to disproportionately contribute to local adaptation due to exposure of recessive alleles to selection, and sex-linked inversions are therefore more likely to sweep to fixation compared to autosomal inversions (103).

For these reasons, X-linkage of this supergene in Sciaridae may have favored its initial emergence as well as its enlargement along the chromosome. Rather than one long paracentric inversion, our analysis suggests the X’ chromosome has undergone multiple rearrangements, which may be explained by multiple adjacent and/or overlapping inversions, smaller inversions nested within larger ones, or some combination thereof, which have accumulated in a stepwise process to suppress recombination along the chromosome. Some of the smaller strata we identified appeared to be far less diverged than others, which may represent gaps between inversion breakpoints. Alternatively, the inversion(s) of the X’ may lead to complex pairing with the X, which may result in varying recombination rates along the chromosome and produce regions of differing divergence. Further resolution of the X’ structure will be required to determine the precise formation of the chromosome. One possibility is that an initial inversion, presumably around the sex-determining region, may have been followed by subsequent adjacent inversions. If true, the accumulation of inversions along the chromosome happened rapidly, as we estimated the ages of all strata as between 0.02 and 0.5 mya. Expansion of the non-recombining region through additional inversions may have adaptively captured female-beneficial alleles at nearby loci, as sex chromosome evolution theory posits (22–24, 104), although this may be hindered by genetic conflict between XX and X’X females.

An alternative explanation for this stratification of the X’ is that sex determination relies on more than one locus, i.e. it is polygenic, and that successive inversions have emerged to control the sex ratio. Among digenic Sciaridae, sex ratios vary significantly: in *Bradysia ocellaris* and *Bradysia matrogrossensis*, broods frequently depart from the expected 50:50 sex ratio and are often heavily skewed in either direction (39, 105). The sex ratio in *B. ocellaris* is heritable, and male-production can evolve from female-production and vice-versa in as few as six generations (106). Taken together, these observations suggest that multiple loci might be involved in sex determination. Monogenic Sciaridae presumably evolved from digenic ancestors, which may have occurred through the adaptive linkage of sex-determining alleles through inversions. Given the young age of the X’, multi-allelic sex determination and the repeated evolution of this phenomenon in Sciaridae may have been favored under certain circumstances over the ancestral digenic sex determination system.

### Evolutionary perspectives on monogenic reproduction in fungus gnats

Within the fungus gnat clade Sciaridae, the relationship between the monogenic and digenic reproductive strategies remains poorly understood, as does the origin of monogenic reproduction. At least one other monogenic species, *B. impatiens*, is known to harbor an X-linked inversion polymorphism. Monogenic reproduction also occurs in many *Bradysia* species, including *B. varians* (10), *B. spatitergum, B. molokaiensis* (35) and *B. paupera* (107), as well as more distantly related Sciaridae including *Scatopsciara nacta* (38), *Ctenosciara hawaiiensis, Lycoriella solispina* (35), *Corynoptera subtrivialis* (10) and *Rhyncosciara hollaenderi* (44). In this respect, our finding that the *B. coprophila* X’ chromosome evolved <0.5 mya has intriguing consequences for understanding the evolution of this reproductive strategy. Unlike the *B. coprophila* X’, the *B. impatiens* X’ non-recombining region is terminal (14), suggesting that the X’ chromosomes in the two species may not be homologous by descent (alternatively, the region has expanded in *B. impatiens* or the terminal portion has re-inverted in *B. coprophila*). Another possibility is that the X’ chromosome in *B. coprophila* may be older than our findings suggest but appears younger due to occasional recombination through gene conversion or double crossovers, which occur within large inversions (108). While crossing-over requires synapsis between chromosomes, gene conversion, i.e. the non-reciprocal copying of stretches of sequence between sister chromatids to repair mismatch errors during replication, does not (109, 110). Furthermore, in *B. impatiens*, dicentric chromatids were observed to form through pairing between the X and X’ along the length of the inversion (14). If such pairing occasionally occurs in *B. coprophila*, it may prevent sequence divergence between the two chromosomes.

Nonetheless, the distribution of monogenic reproduction among sciarids indicates multiple evolutionary origins. For example, within *Bradysia* alone, both monogenic and digenic species exist (107), and the same pattern is found in more distantly related genera (35). If sex determination involves multiple loci, inversions may have emerged in some lineages to fix the production of sex-biased broods in a particular direction. However, this raises the question: what drives the turnover between reproductive strategies in this clade? Haig (111) suggested that female production evolved as a response to a male-biased sex ratio. Fungus gnats carry a unique type of chromosome that are only found in the germline (germline-restricted chromosomes, GRCs), in addition to their sex chromosomes and autosomes. The GRCs are disproportionately transmitted by males, and so may have distorted the sex ratio in their favour (111). Presence of GRCs in Sciaridae does appear to correlate with monogenic reproduction, and many (but not all) digenic species lack them (112, 113). In support of this is the observation that a monogenic lab-reared line of *B. impatiens* reverted to digenic reproduction following loss of its GRCs (114), though similar findings have not yet been repeated. However, the potential role of the GRCs in sex determination in Sciaridae remains unknown. Interestingly, the GRCs of *B. coprophila* were recently found to have introgressed into Sciaridae following an ancient hybridization event with Cecidomyiidae (47), a clade that shares many features with Sciaridae including paternal genome elimination, GRCs, sex determination by chromosome elimination, and monogenic reproduction (9, 16). It is thus tempting to speculate that GRCs may have spread through Sciaridae via similar introgression events, and that this may have also facilitated the spread of monogenic reproduction. If female-determining inversions were to repeatedly evolve, individuals lacking inversions would be selected to increase their male production as an evolutionary response, with the expected result being that the X’X genotype is maintained at 50% in the population by frequency-dependent selection. However, the fact that we observe X’ degeneration could mean that X’X females will, over time, have reduced fitness, which should favor the invasion of individuals capable of digenic reproduction, unless selective forces or rare recombination exists to keep the X’ from degrading significantly. Future work on the role of GRCs in sciarid sex determination, and on the relationship between the unusual genetic aspects across different sciarid species, will be required to elucidate the origins and turnover of sex determinations strategies in this clade

## Materials and Methods

### Data collection

The *Bradysia* (formerly *Sciara*) *coprophila* strain used in this study was obtained from the *Sciara* stock center at Brown University (https://sites.brown.edu/sciara/), which is derived from highly inbred individuals reared in laboratory conditions since the 1920s (51). Data used in this study was produced at both Edinburgh University and the Carnegie Institution for Science in Baltimore. At Edinburgh, DNA was extracted using a version of the Qiagen DNeasy Blood and Tissue Kit extraction protocol modified for high-molecular weight (HMW) extractions. DNA from 50-60 pooled X’X heads was used for sequencing on the Illumina NovaSeq S1 platform for paired-end 150bp reads with 350bp inserts; DNA from 30-40 X’X heads was used for PacBio Single-Molecular, Real-Time long-read sequence data at 50x coverage. All DNA samples were quantified using the Qubit (ThermoFisher). HiC data was sequenced from approximately 50 pooled whole X’X females, which were ground using a DiagoCine Powermasher II with a Biomasher II attachment; libraries were prepared and sequenced for around 330x coverage by Science for Life Laboratory in Stockholm, Sweden. At Carnegie, DNA was extracted using DNAzol (ThermoFisher) from 20-36 pooled whole-body individuals, two replicates per genotype (X’X, XX, X0), quantified with the Qubit (ThermoFisher), analyzed for purity with Nanodrop (ThermoFisher), analyzed for HMW integrity with 0.5% agarose gel electrophoresis, prepared for sequencing using Illumina Nextera reagents, and was sequenced on the Illumina NextSeq platform to generate 75bp paired-end reads of ∼150-400bp fragments. X0 PacBio data was from male embryos, and was generated previously as part of assembling the original somatic reference genome (42). X0 HiC data was from male pupae, and was generated previously for chromosome-scale scaffolding of the somatic reference genome (43). Illumina and HiC reads were adapter- and quality-trimmed using fastp v0.2.1 (115) and quality was assessed before and after trimming using FastQC (116).

### Analysis of X-X’ divergence and evolutionary strata

To identify SVs, 75bp paired-end Illumina reads and PacBio reads from X’X and X0 samples were aligned to the X0 reference genome (43) with BWA-MEM (117) to force-map X’ reads to the X chromosome. SAMtools (118) was used to sort, merge and index BAM files prior to calling SVs from Illumina alignments with Smoove v0.2.8 (119, 120) and PacBio alignments with Sniffles v2.0 (121, 122). svtools (123) was used to convert variant files to bedpe files. To target fixed variants between X and X’ and thus exclude stochastic variants present only in one or some individuals, at least 4 reads were required to support a variant call. The R/Bioconductor (124) package Sushi (125) was used to plot SVs. HiC reads were aligned to the reference genome using Juicer (126), and HiC contact heatmaps were produced using the script HiC_view.py (https://github.com/A-J-F-Mackintosh/Mackintosh_et_al_2022_Ipod, 127). To call SNVs, the Illumina alignments were processed with Picardtools (128) before calling variants with the GATK-4 best practices pipeline (129, 130). The python scripts parseVCF.py and popgenWindows.py (https://github.com/simonhmartin/genomics_general) were then used to parse the variant files and calculate the density of SNVs across 100 Kb windows, respectively. The R package bcp (131– 133), which employs a Bayesian approach to identify change points between clusters of means, was used to identify putative evolutionary strata and to delineate the X region homologous to the inversion, and the R (124) package ggplot2 (134) was used to plot SNVs, posterior means and point changes.

Two methods were employed to estimate the ages of strata. Here we assumed a neutral mutation rate and a similar mutation rate to *Drosophila melanogaster*, i.e. between 2.8 × 10^−9^ (135) and 4.9 × 10^−9^ (136), and a 24-40 day generation time for *B. coprophila* (**S4 Text**). First, after mapping X’X reads to the X0 genome (see above), heterozygous SNPs and invariant (homozygous) sites were counted along the 58.8Mb X region homologous to the inversion, and *Dxy* was calculated as the total number of heterozygous sites divided by the total number of homozygous and heterozygous sites. Estimates for the number of generations since divergence were then calculated as “*Dxy*/2*r”, where r is mutation rate. Second, we aimed to produce a more accurate estimate by targeting only synonymous (neutral) mutations. SnpEff (137) and SnpSift (138) were used to annotate variants and count the number of synonymous variants per gene. The script partitionCDS.py (https://github.com/A-J-F-Mackintosh/Mackintosh_et_al_2022_Ipod, 127) was used to annotate degeneracy for all genic sites, and we subsequently calculated the number of synonymous sites per gene for single-copy X-X’ homologs (see below) within each stratum. Divergence in generations for each gene was then calculated as “V/(2*r*S)”, where V is the number of synonymous variants, r is mutation rate, and S is the number of synonymous sites.

### Genome assembly and annotation

A genome assembly was initially generated *de* novo from PacBio reads from X’X females, but only around 3.6Mb of sequence from this assembly could be assigned to the X’ inversion because high sequence similarity between X and X’ chromosomes and high read error rates (∼11-16% on average) caused them to collapse upon assembly (**S1 Text, Figure S1**). To assemble the X’ inversion, putative X’-specific 27-mers were identified using KAT (139), counted using KMC3 (140) and fasta files of 27-mers were obtained using custom python scripts (47, https://github.com/RossLab/Bradysia-GRCs/). 27-mers with a frequency of 0 in male reads and a frequency of greater than 5 in X’X female reads (to exclude 27-mers containing read errors) were used to identify reads originating from the X’ inversion. Cookiecutter (141) was used to parse inversion and non-inversion (A + X) reads from trimmed X’X female Illumina paired-end 150bp and 75bp read files, whereby a read is extracted from the raw file if it contains a k-mer from a given k-mer library. The X’ inversion-specific reads were assembled using SPAdes (142) to yield 61.77 Mb spanning 42564 contigs. These contigs were scaffolded using RagTag (143) and the X chromosome scaffold as a reference (43) to produce a 52.43 Mb assembly containing 9409 gaps; contigs that failed to scaffold were discarded. The inversion scaffold was then combined with the male X0 genome to produce an X’X genome. PacBio long reads from an X’X female were mapped to the X’X genome with minimap2 and gaps were plugged using Racon (144). Illumina reads from an X’X female were subsequently mapped to the X’X genome with minimap2 (145) and Racon (144) was used to polish the assembly. After gap-plugging and polishing the X’ inversion scaffold totalled 55.09 Mb and contained 2790 gaps.

The genome was soft-masked prior to annotation. To this end, *de novo* repeat families were created using RepeatModeler v2.0 (146), which were then combined with known dipteran repeat families from RepBase (147). RepeatMasker v4.0 (148) was used to soft-mask the genome; in total 36.36% of bases were soft-masked. The genome was then annotated using BRAKER2 (149–153) with RNAseq alignments and homology-based datasets (**S5 Text**). Functional information for the 26887 protein sequences in the resulting BRAKER2 gene annotation set was then obtained by finding the best BLASTP (154) hits in several protein databases and with InterProScan (155, **S5 Text**). To annotate transposable elements (TEs) in the genome, the module reasonaTE of the TranposonUltimate v1.03 pipeline (166) was used (**S6 Text**).

### Analysis of functional degradation

We identified X-X’ homologous gene copies by running OrthoFinder (156) on the longest predicted transcripts from each chromosome. To identify disrupted X’-linked genes, genotyped variant files for X’X and X0 75bp paired-end Illumina reads mapped against the Bcop_v2 (X, II, III, IV) genome (as described above) were filtered for SNVs and indels using SnpEff (137), and SnpSift (138) was used to extract variant information. SNV and indel counts were then analyzed in RStudio (124) and only genes from the identified 2321 single-copy homologs were considered. Two replicates of each genotype (X’X and X0) were aligned, and only consensus variant calls between two replicates were considered to identify fixed differences between X and X’ and exclude low-frequency variants present in only one or a few individuals. We identified variants specific to the X’ by excluding common variants also found with X0 male data, which may represent polymorphisms on the X or errors in the X chromosome reference sequence (43). Genes with loss or gain of start or stop codons or indels causing frameshifts were counted as disrupted.

To identify silenced genes, we parsed X and X’ RNAseq reads using elements of a pipeline developed for (157, https://github.com/MooHoll/Parent_of_Origin_Expression_Bumblebee) to avoid mismapping between the two chromosomes. To this end, we created an N-masked version of the X chromosome to mask SNVs between the X and X’ by mapping reads from X’X individuals to the X0 reference genome (43) using Bowtie2 (158), processed BAM files using Picardtools (128) and SAMtools (118) and called and filtered SNVs using freebayes (159) and VCFtools (160) respectively. BEDtools (161) was used to identify X- and X’-specific variants and to N-mask those variants on the X chromosome prior to mapping RNAseq reads. We mapped RNAseq reads from pooled female data from four life stages: embryo, larva, pupa and adult (42) to the N-masked genome using STAR (162) and subsequently assigned RNAseq reads to the X or X’ chromosome using SNPsplit (163). Gene expression was quantified using Kallisto (164) with the 2321 single-copy homologs we identified as a Kallisto index, and counts were normalized using EdgeR (165). Genes with zero counts of TPM (transcripts per million) were assumed to be non-expressed. We also included genes with TPMs in the bottom 0.1% of non-zero TPM counts within a sample as non-expressed to account for stochastic mismapping of RNAseq reads. We counted X’ genes as ‘silenced’ if the X copy was expressed but the X’ copy was not, and if genes were both silenced and contained pseudogenizing mutations we counted them as ‘silenced and disrupted’. Finally, analysis and plotting of counts was carried out with R Studio (124) with use of the ggplot2 package (134).

To examine DC of X genes that have corresponding degraded X’ homologs, we applied the same pipeline described above to pull X-specific reads (separated from X’-specific reads) from X’X and XX samples (**S7 Text**). Kallisto (164) was then used to quantify the resulting X reads against the 2321 X-linked single-copy homologs. After normalization between the samples with EdgeR (165). We assumed that in X’X females, X-linked genes with degraded X’ homologs should have upregulated expression if they are dosage compensated, so as to match expression in XX females. We therefore identified upregaulted genes as those with a log2FC (fold-change) of greater than 0.5.

## Acknowledgments

The authors thank Allan C. Spradling, an investigator of the Howard Hughes Medical Institute (HHMI), for providing funding for data generated by JMU. The authors acknowledge support from the National Genomics Infrastructure in Stockholm funded by Science for Life Laboratory, the Knut and Alice Wallenberg Foundation and the Swedish Research Council, and SNIC/Uppsala Multidisciplinary Center for Advanced Computational Science for assistance with massively parallel sequencing and access to the UPPMAX computational infrastructure. We thank Allison Pinder and Frederick Tan from the Carnegie Sequencing Core for operating the Illumina NextSeq and demultiplexing the data. Some of the computations presented here were conducted through Carnegie’s partnership in the Resnick High Performance Computing Centre, a facility supported by Resnick Sustainability Institute at the California Institute of Technology. RBB, CNH, KSJ, AJM and LR were funded by a European Research Council Starting Grant (PGErepro, to LR). MG was funded by the Novo Nordisk Foundation (NNF18OC0030954) We thank Deborah Charlesworth for providing comments on the manuscript. We also thank all present and past members of Ross Lab for useful discussion throughout this project.

## Data availability

Data used in this study will become available upon publication.

## References

1. L. W. Beukeboom, N. Perrin, The evolution of sex determination (Oxford University Press, USA, 2014).

2. P. A. Saunders, F. Veyrunes, Unusual Mammalian Sex Determination Systems: A Cabinet of Curiosities. Genes 12, 1770 (2021).

3. D. Bachtrog, et al., Sex determination: why so many ways of doing it? PLoS Biol. 12, e1001899 (2014).

4. J. J. Bull, The Evolution of Sex Determining Mechanisms (Benjamin Cummings, 1983).

5. D. N. Roy, L. B. Siddons, On the life history and bionomics of Chrysomyia rufifacies Macq. (Order Diptera, Family Calliphoridae). Parasitology 31, 442–447 (1939).

6. F.-H. Ullerich, Monogene Fortpflanzung bei der Fliege Chrysomyia albiceps. Z. Für Naturforschung B 13, 473–474 (1958).

7. I. Geyer-Duszyńska, Spindle disappearance and chromosome behavior after partial-embryo irradiation in Cecidomyiidae (Diptera). Chromosoma 12, 233–247 (1961).

8. Z. Kraczkiewicz, Premiers stades de l’oogenèse de Rhabdophaga saliciperda (Cecidomyiidae, Diptera). Chromosoma 18, 208–229 (1966).

9. J. J. Stuart, J. H. Hatchett, Cytogenetics of the Hessian Fly: I. Mitotic Karyotype Analysis and Polytene Chromosome Correlations. J. Hered. 79, 184–189 (1988).

10. C. W. Metz, Chromosome behavior, inheritance and sex determination in Sciara. Am. Nat. 72, 485–520 (1938).

11. S. A. Gerbi, “Unusual Chromosome Movements in Sciarid Flies” in Germ Line — Soma Differentiation, Results and Problems in Cell Differentiation., W. Hennig, Ed. (Springer, 1986), pp. 71–104.

12. L. Sánchez, Sciara as an experimental model for studies on the evolutionary relationships between the zygotic, maternal and environmental primary signals for sexual development. J. Genet. 89, 325–331 (2010).

13. M. J. Scott, M. L. Pimsler, A. M. Tarone, Sex Determination Mechanisms in the Calliphoridae (Blow Flies). Sex. Dev. 8, 29–37 (2014).

14. H. L. Carson, The selective elimination of inversion dicentric chromatids during meiosis in the eggs of Sciara impatiens. Genetics 31, 95–113 (1946).

15. H. V. Crouse, X heterochromatin subdivision and cytogenetic analysis in Sciara coprophila (diptera, sciaridae) - II. The controlling element. Chromosoma 74, 219–239 (1979).

16. T. R. Benatti, et al., A Neo-Sex Chromosome That Drives Postzygotic Sex Determination in the Hessian Fly (Mayetiola destructor). Genetics 184, 769–777 (2010).

17. B. Vicoso, Molecular and evolutionary dynamics of animal sex-chromosome turnover. Nat. Ecol. Evol. 3, 1632–1641 (2019).

18. A. E. Wright, et al., Convergent recombination suppression suggests role of sexual selection in guppy sex chromosome formation. Nat. Commun. 8, 14251 (2017).

19. T. Connallon, et al., Local adaptation and the evolution of inversions on sex chromosomes and autosomes. Philos. Trans. R. Soc. B Biol. Sci. 373, 20170423 (2018).

20. T. Lenormand, D. Roze, Y recombination arrest and degeneration in the absence of sexual dimorphism. Science 375, 663–666 (2022).

21. P. Jay, E. Tezenas, A. Véber, T. Giraud, Modeling the stepwise extension of recombination suppression on sex chromosomes and other supergenes through deleterious mutation sheltering. 2021.05.17.444504 (2022).

22. D. Charlesworth, B. Charlesworth, Sex differences in fitness and selection for centric fusions between sex-chromosomes and autosomes. Genet. Res. 35, 205–214 (1980).

23. W. R. Rice, The Accumulation of Sexually Antagonistic Genes as a Selective Agent Promoting the Evolution of Reduced Recombination between Primitive Sex Chromosomes. Evolution 41, 911–914 (1987).

24. D. Charlesworth, B. Charlesworth, G. Marais, Steps in the evolution of heteromorphic sex chromosomes. Heredity 95, 118–128 (2005).

25. W. G. Hill, A. Robertson, The effect of linkage on limits to artificial selection. Genet. Res. 8, 269–294 (1966).

26. J. Felsenstein, The evolutionary advantage of recombination. Genetics 78, 737–756 (1974).

27. D. Bachtrog, E. Hom, K. M. Wong, X. Maside, P. de Jong, Genomic degradation of a young Y chromosome in Drosophila miranda. Genome Biol. 9, R30 (2008).

28. R. Erlandsson, J. F. Wilson, S. Pääbo, Sex Chromosomal Transposable Element Accumulation and Male-Driven Substitutional Evolution in Humans. Mol. Biol. Evol. 17, 804–812 (2000).

29. D. Chalopin, J.-N. Volff, D. Galiana, J. L. Anderson, M. Schartl, Transposable elements and early evolution of sex chromosomes in fish. Chromosome Res. 23, 545–560 (2015).

30. S. Ponnikas, H. Sigeman, J. K. Abbott, B. Hansson, Why Do Sex Chromosomes Stop Recombining? Trends Genet. 34, 492–503 (2018).

31. M. S. Moses, Chas. W. Metz, Evidence that the Female Is Responsible for the Sex Ratio in Sciara (Diptera). Proc. Natl. Acad. Sci. 14, 928–930 (1928).

32. A. M. DuBois, Chromosome behavior during cleavage in the eggs of Sciara coprophila (Diptera) in the relation to the problem of sex determination. Z. Für Zellforsch. Mikrosk. Anat. 19, 595–614 (1933).

33. Chas. W. Metz, M. L. Schmuck, Unisexual progenies and the sex chromosome mechanism in Sciara. Proc. Natl. Acad. Sci. 15, 863–866 (1929).

34. C. W. Metz, Chromosome Behavior, Inheritance and Sex Determination in Sciara. Am. Nat. 72, 485–520 (1938).

35. W. A. Steffan, Laboratory studies and ecological notes on Hawaiian Sciaridae (Diptera). Pac. Insects 16, 41–50 (1974).

36. H. V. Crouse, The nature of the influence of X-translocations on sex of progeny in Sciara coprophila. Chromosoma 11, 146–166 (1960).

37. M. D. McCarthy, Chromosome Studies on Eight Species of Sciara (Diptera) with Special Reference to Chromosome Changes of Evolutionary Significance. Am. Nat. 79, 104–121 (1945).

38. M. D. McCarthy, Chromosome Studies on Eight Species of Sciara (Diptera) with Special Reference to Chromosome Changes of Evolutionary Significance. II (Continued). Am. Nat. 79, 228–245 (1945).

39. L. S. Rocha, A. L. P. Perondini, Analysis of the sex ratio in Bradysia matogrossensis (Diptera, Sciaridae). Genet. Mol. Biol. 23, 97–103 (2000).

40. H. V. Crouse, Translocations in Sciara: their bearing on chromosome behavior and sex determination. Univ Mo. Res Bull 379, 1–75 (1943).

41. H. V. Crouse, X heterochromatin subdivision and cytogenetic analysis in Sciara coprophila (Diptera, Sciaridae) - I. Centromere localization. Chromosoma 63, 39–55 (1977).

42. J. M. Urban, et al., High contiguity de novo genome assembly and DNA modification analyses for the fungus fly, Sciara coprophila, using single-molecule sequencing. BMC Genomics 22, 643 (2021).

43. J. M. Urban, S. A. Gerbi, A. C. Spradling, Chromosome-scale scaffolding of the fungus gnat genome (Diptera: Bradysia coprophila). bioRxiv (2022).

44. E. Mattingly, J. N. Dumont, Early spermatogenesis in Rhynchosciara. In Vitro 6, 286–299 (1971).

45. N. Gabrusewycz-Garcia, Cytological and autoradiographic studies in Sciara coprophila salivary gland chromosomes. Chromosoma 15, 312–344 (1964).

46. E. M. Rasch, Genome size and determination of DNA content of the X chromosomes, autosomes, and germ line-limited chromosomes of Sciara coprophila. J. Morphol. 267, 1316–1325 (2006).

47. C. N. Hodson, K. S. Jaron, S. Gerbi, L. Ross, Gene-rich germline-restricted chromosomes in black-winged fungus gnats evolved through hybridization. PLOS Biol. 20, e3001559 (2022).

48. D. H. Palmer, T. F. Rogers, R. Dean, A. E. Wright, How to identify sex chromosomes and their turnover. Mol. Ecol. 28, 4709–4724 (2019).

49. S. Koren, et al., De novo assembly of haplotype-resolved genomes with trio binning. Nat. Biotechnol. 36, 1174–1182 (2018).

50. C. W. Metz, Genetic Evidence of a Selective Segregation of Chromosomes in Sciara (Diptera). Proc. Natl. Acad. Sci. 12, 690–692 (1926).

51. Chas. W. Metz, Chromosomes and Sex in Sciara. Science 61, 212–214 (1925).

52. S. Zaffran, M. Astier, D. Gratecos, M. Sémériva, The held out wings (how) Drosophila gene encodes a putative RNA-binding protein involved in the control of muscular and cardiac activity. 12.

53. P. N. Adler, “The frizzled/stan Pathway and Planar Cell Polarity in the Drosophila Wing” in Current Topics in Developmental Biology, (Elsevier, 2012), pp. 1–31.

54. B. Vicoso, D. Bachtrog, Numerous Transitions of Sex Chromosomes in Diptera. PLoS Biol. 13, e1002078 (2015).

55. H. Blackmon, J. P. Demuth, Estimating Tempo and Mode of Y Chromosome Turnover: Explaining Y Chromosome Loss With the Fragile Y Hypothesis. Genetics 197, 561–572 (2014).

56. L. Z. Carabajal Paladino, et al., Sex Chromosome Turnover in Moths of the Diverse Superfamily Gelechioidea. Genome Biol. Evol. 11, 1307–1319 (2019).

57. M. Nozawa, et al., Shared evolutionary trajectories of three independent neo-sex chromosomes in Drosophila. Genome Res. 31, 2069–2079 (2021).

58. M. Dalíková, et al., New Insights into the Evolution of the W Chromosome in Lepidoptera. J. Hered. 108, 709–719 (2017).

59. H. J. Muller, The Relation of Recombination to Mutational Advance. Mutat. Res. 1, 2–9 (1964).

60. J. Maynard-Smith, J. Haigh, The hitch-hiking effect of a favourable gene. Genet. Res. 23, 23–35 (1974).

61. B. Charlesworth, The evolution of sex chromosomes. Science 251, 1030–1033 (1991).

62. J. Engelstadter, Constraints on the evolution of asexual reproduction. Bioessays 30, 1138–1150 (2008).

63. A. B. Carvalho, A. G. Clark, Y Chromosome of D. pseudoobscura Is Not Homologous to the Ancestral Drosophila Y. Science 307, 108–110 (2005).

64. S. Mahajan, D. Bachtrog, Convergent evolution of Y chromosome gene content in flies. Nat. Commun. 8, 785 (2017).

65. D. Bachtrog, B. Charlesworth, Reduced adaptation of a non-recombining neo-Y chromosome. Nature 416, 323–326 (2002).

66. M. Steinemann, S. Steinemann, “Enigma of Y chromosome degeneration: Neo-Y and Neo-X chromosomes of Drosophila miranda a model for sex chromosome evolution” in Mutation and Evolution, Contemporary Issues in Genetics and Evolution., R. C. Woodruff, J. N. Thompson, Eds. (Springer Netherlands, 1998), pp. 409–420.

67. D. Bachtrog, Expression Profile of a Degenerating Neo-Y Chromosome in Drosophila. Curr. Biol. 16, 1694–1699 (2006).

68. S. Mahajan, K. H.-C. Wei, M. J. Nalley, L. Gibilisco, D. Bachtrog, De novo assembly of a young Drosophila Y chromosome using single-molecule sequencing and chromatin conformation capture. PLOS Biol. 16, e2006348 (2018).

69. C. P. Vieira, P. A. Coelho, J. Vieira, Inferences on the Evolutionary History of the Drosophila americana Polymorphic X / 4 Fusion From Patterns of Polymorphism at the X -Linked paralytic and elav Genes. Genetics 164, 1459–1469 (2003).

70. Q. Zhou, D. Bachtrog, Ancestral Chromatin Configuration Constrains Chromatin Evolution on Differentiating Sex Chromosomes in Drosophila. PLOS Genet. 11, e1005331 (2015).

71. K. H.-C. Wei, D. Bachtrog, Ancestral male recombination in Drosophila albomicans produced geographically restricted neo-Y chromosome haplotypes varying in age and onset of decay. PLOS Genet. 15, e1008502 (2019).

72. B. Charlesworth, J. A. Coyne, N. H. Barton, The Relative Rates of Evolution of Sex Chromosomes and Autosomes. Am. Nat. 130, 113–146 (1987).

73. S. Ohno, Sex Chromosomes and Sex-Linked Genes (Springer Science & Business Media, 1967).

74. S. R. Schulze, L. L. Wallrath, Gene Regulation by Chromatin Structure: Paradigms Established in Drosophila melanogaster. Annu. Rev. Entomol. 52, 171–192 (2007).

75. S. V. Flores, A. L. Evans, B. F. McAllister, Independent Origins of New Sex-Linked Chromosomes in the melanica and robusta Species Groups of Drosophila. BMC Evol. Biol. 8, 33 (2008).

76. C. E. Ellison, D. Bachtrog, Dosage Compensation via Transposable Element Mediated Rewiring of a Regulatory Network. Science 342, 846–850 (2013).

77. I. Marín, A. Franke, G. J. Bashaw, B. S. Baker, The dosage compensation system of Drosophila is co-opted by newly evolved X chromosomes. Nature 383, 160–163 (1996).

78. Q. Zhou, et al., The Epigenome of Evolving Drosophila Neo-Sex Chromosomes: Dosage Compensation and Heterochromatin Formation. PLoS Biol. 11, e1001711 (2013).

79. M. Nozawa, K. Ikeo, T. Gojobori, Gene-by-gene or localized dosage compensation on the neo-X chromosome in Drosophila miranda. Genome Biol. Evol. (2018).

80. P. R. da Cunha, B. Granadino, A. L. Perondini, L. Sánchez, Dosage compensation in sciarids is achieved by hypertranscription of the single X chromosome in males. Genetics 138, 787–790 (1994).

81. Q. Zhou, D. Bachtrog, Sex-Specific Adaptation Drives Early Sex Chromosome Evolution in Drosophila. Science 337, 341–345 (2012).

82. C. L. Peichel, et al., Assembly of the threespine stickleback Y chromosome reveals convergent signatures of sex chromosome evolution. Genome Biol. 21, 177 (2020).

83. M. Petersen, et al., Diversity and evolution of the transposable element repertoire in arthropods with particular reference to insects. BMC Ecol. Evol. 19, 11 (2019).

84. V. Peona, et al., The avian W chromosome is a refugium for endogenous retroviruses with likely effects on female-biased mutational load and genetic incompatibilities. Philos. Trans. R. Soc. B Biol. Sci. 376, 20200186 (2021).

85. Y.-J. Kim, J. Lee, K. Han, Transposable Elements: No More “Junk DNA.” Genomics Inform. 10, 226 (2012).

86. M. Joron, et al., Chromosomal rearrangements maintain a polymorphic supergene controlling butterfly mimicry. Nature 477, 203–206 (2011).

87. W. Zhang, E. Westerman, E. Nitzany, S. Palmer, M. R. Kronforst, Tracing the origin and evolution of supergene mimicry in butterflies. Nat. Commun. 8, 1269 (2017).

88. P. Jay, et al., Supergene Evolution Triggered by the Introgression of a Chromosomal Inversion. Curr. Biol. 28, 1839-1845.e3 (2018).

89. A. J. Mongue, A. Y. Kawahara, Population differentiation and structural variation in the Manduca sexta genome across the United States. G3 GenesGenomesGenetics 12, jkac047 (2022).

90. A. J. Mongue, M. E. Hansen, J. R. Walters, Support for faster and more adaptive Z chromosome evolution in two divergent lepidopteran lineages*. Evolution 76, 332–345 (2022).

91. J. Wang, et al., A Y-like social chromosome causes alternative colony organization in fire ants. Nature 493, 664–668 (2013).

92. Z. Yan, et al., Evolution of a supergene that regulates a trans-species social polymorphism. Nat. Ecol. Evol. 4, 240–249 (2020).

93. S. Branco, et al., Multiple convergent supergene evolution events in mating-type chromosomes. Nat. Commun. 9, 2000 (2018).

94. E. R. Funk, et al., A supergene underlies linked variation in color and morphology in a Holarctic songbird. Nat. Commun. 12, 6833 (2021).

95. C. Küpper, et al., A supergene determines highly divergent male reproductive morphs in the ruff. Nat. Genet. 48, 79–83 (2016).

96. K.-W. Kim, et al., A sex-linked supergene controls sperm morphology and swimming speed in a songbird. Nat. Ecol. Evol. 1, 1168–1176 (2017).

97. T. Schwander, R. Libbrecht, L. Keller, Supergenes and Complex Phenotypes. Curr. Biol. 24, R288–R294 (2014).

98. M. J. Thompson, C. D. Jiggins, Supergenes and their role in evolution. Heredity 113, 1–8 (2014).

99. J. A. Coyne, Genetics and speciation. Nature 355, 511–515 (1992).

100. D. C. Presgraves, Sex chromosomes and speciation in Drosophila. Trends Genet. 24, 336–343 (2008).

101. E. M. Tuttle, et al., Divergence and Functional Degradation of a Sex Chromosome-like Supergene. Curr. Biol. 26, 344–350 (2016).

102. K.-W. Kim, et al., Stepwise evolution of a butterfly supergene via duplication and inversion. 377, 20210207 (2022).

103. C. Lasne, C. M. Sgrò, T. Connallon, The Relative Contributions of the X Chromosome and Autosomes to Local Adaptation. Genetics 205, 1285–1304 (2017).

104. H. Blackmon, Y. Brandvain, Long-Term Fragility of Y Chromosomes Is Dominated by Short-Term Resolution of Sexual Antagonism. Genetics 207, 1621–1629 (2017).

105. R. G. Nigro, M. C. C. Campos, A. L. P. Perondini, Temperature and the progeny sex-ratio in Sciara ocellaris (Diptera, Sciaridae). Genet. Mol. Biol. 30, 152–158 (2007).

106. B. Davidheiser, Observations on the Inheritance of Sex in Sciara Ocellaris (Diptera). Ohio J. Sci. 47, 89–102 (1947).

107. Y. Liu, “Chemoecological Studies on the Reproductive Behaviors of the Darkwinged Fungus Gnat, Bradysia paupera (Diptera: Sciaridae, doctoral thesis),” University of Tsukuba. (2007).

108. A. Navarro, E. Betrán, A. Barbadilla, A. Ruiz, Recombination and Gene Flux Caused by Gene Conversion and Crossing Over in Inversion Heterokaryotypes. Genetics 146, 695–709 (1997).

109. J. W. Szostak, T. L. Orr-Weaver, R. J. Rothstein, F. W. Stahl, The double-strand-break repair model for recombination. Cell 33, 25–35 (1983).

110. M. S. McMahill, C. W. Sham, D. K. Bishop, Synthesis-Dependent Strand Annealing in Meiosis. PLoS Biol. 5, e299 (2007).

111. D. Haig, The evolution of unusual chromosomal systems in sciarid flies: intragenomic conflict and the sex ratio. J. Evol. Biol. 6, 249–261 (1993).

112. C. N. Hodson, L. Ross, Evolutionary Perspectives on Germline-Restricted Chromosomes in Flies (Diptera). Genome Biol. Evol. 13, evab072 (2021).

113. S. A. Gerbi, Non-random chromosome segregation and chromosome eliminations in the fly Bradysia (Sciara). Chromosome Res. (2022).

114. H. V. Crouse, A. Brown, B. C. Mumford, L-chromosome inheritance and the problem of chromosome “imprinting” in Sciara (Sciaridae, Diptera). Chromosoma 34, 324–339 (1971).

115. S. Chen, Y. Zhou, Y. Chen, J. Gu, fastp: an ultra-fast all-in-one FASTQ preprocessor. bioRxiv, 274100–274100 (2018).

116. S. Andrews, others, FastQC: a quality control tool for high throughput sequence data (2010).

117. H. Li, Aligning sequence reads, clone sequences and assembly contigs with BWA-MEM. ArXiv Prepr. ArXiv13033997 (2013).

118. H. Li, et al., The Sequence Alignment/Map format and SAMtools. Bioinformatics 25, 2078–2079 (2009).

119. B. S. Pedersen, R. Layer, A. Quilan, smoove: structural-variant calling and genotyping with existing tools (2020).

120. R. M. Layer, C. Chiang, A. R. Quinlan, I. M. Hall, LUMPY: a probabilistic framework for structural variant discovery. Genome Biol. 15, R84 (2014).

121. F. J. Sedlazeck, et al., Accurate detection of complex structural variations using single-molecule sequencing. Nat. Methods 15, 461–468 (2018).

122. M. Smolka, et al., “Comprehensive Structural Variant Detection: From Mosaic to Population-Level” (Bioinformatics, 2022).

123. D. E. Larson, et al., svtools: population-scale analysis of structural variation. Bioinformatics 35, 4782–4787 (2019).

124. R Core Team, R: A Language and Environment for Statistical Computing (R Foundation for Statistical Computing, 2022).

125. D. H. Phanstiel, A. P. Boyle, C. L. Araya, M. P. Snyder, Sushi.R: flexible, quantitative and integrative genomic visualizations for publication-quality multi-panel figures. Bioinformatics 30, 2808–2810 (2014).

126. N. C. Durand, et al., Juicer Provides a One-Click System for Analyzing Loop-Resolution Hi-C Experiments. Cell Syst. 3, 95–98 (2016).

127. A. Mackintosh, et al., The genome sequence of the scarce swallowtail, Iphiclides podalirius. G3 GenesGenomesGenetics 12, jkac193 (2022).

128. , Picard toolkit. Broad Inst. GitHub Repos. (2019).

129. A. McKenna, et al., The Genome Analysis Toolkit: a MapReduce framework for analyzing next-generation DNA sequencing data. Genome Res. (2010).

130. M. A. DePristo, et al., A framework for variation discovery and genotyping using next-generation DNA sequencing data. Nat. Genet. 43, 491–498 (2011).

131. C. Erdman, J. W. Emerson, bcp: an R package for performing a Bayesian analysis of change point problems. J. Stat. Softw. 23, 1–13 (2008).

132. C. Erdman, J. W. Emerson, bcp: A Package for Performing a Bayesian Analysis of Change Point Problems. R package version 1.8. 4, URL http://CRAN.R-project.org (2007).

133. X. Wang, J. W. Emerson, Bayesian change point analysis of linear models on graphs. ArXiv Prepr. ArXiv150900817 (2015).

134. H. Wickham, “Data analysis” in Ggplot2, (Springer, 2016), pp. 189–201.

135. P. D. Keightley, R. W. Ness, D. L. Halligan, P. R. Haddrill, Estimation of the Spontaneous Mutation Rate per Nucleotide Site in a Drosophila melanogaster Full-Sib Family. Genetics 196, 313–320 (2014).

136. Z. J. Assaf, S. Tilk, J. Park, M. L. Siegal, D. A. Petrov, Deep sequencing of natural and experimental populations of Drosophila melanogaster reveals biases in the spectrum of new mutations. Genome Res. 27, 1988–2000 (2017).

137. P. Cingolani, et al., A program for annotating and predicting the effects of single nucleotide polymorphisms, SnpEff. Fly (Austin) 6, 80–92 (2012).

138. P. Cingolani, et al., Using Drosophila melanogaster as a Model for Genotoxic Chemical Mutational Studies with a New Program, SnpSift. Front. Genet. 3 (2012).

139. D. Mapleson, G. Garcia Accinelli, G. Kettleborough, J. Wright, B. J. Clavijo, KAT: a K-mer analysis toolkit to quality control NGS datasets and genome assemblies. Bioinformatics, btw663 (2016).

140. M. Kokot, M. Długosz, S. Deorowicz, KMC 3: counting and manipulating k-mer statistics. Bioinformatics 33, 2759–2761 (2017).

141. E. Starostina, G. Tamazian, P. Dobrynin, S. O’Brien, A. Komissarov, “Cookiecutter: a tool for kmer-based read filtering and extraction” Bioinformatics (2015).

142. A. Bankevich, et al., SPAdes: a new genome assembly algorithm and its applications to single-cell sequencing. J. Comput. Biol. 19, 455–477 (2012).

143. M. Alonge, et al., “Automated assembly scaffolding elevates a new tomato system for high-throughput genome editing” Plant Biology (2021).

144. R. Vaser, I. Sović, N. Nagarajan, M. Šikić, Fast and accurate de novo genome assembly from long uncorrected reads. Genome Res. 27, 737–746 (2017).

145. H. Li, Minimap2: pairwise alignment for nucleotide sequences. Bioinformatics 34, 3094–3100 (2018).

146. J. M. Flynn, et al., RepeatModeler2 for automated genomic discovery of transposable element families. Proc. Natl. Acad. Sci. 117, 9451–9457 (2020).

147. W. Bao, K. K. Kojima, O. Kohany, Repbase Update, a database of repetitive elements in eukaryotic genomes. Mob. Dna 6, 1–6 (2015).

148. A. Smit, R. Hubley, P. Green, RepeatMasker Open-4.0. 2013–2015 (2015).

149. M. Stanke, M. Diekhans, R. Baertsch, D. Haussler, Using native and syntenically mapped cDNA alignments to improve de novo gene finding. Bioinformatics 24, 637–644 (2008).

150. K. J. Hoff, S. Lange, A. Lomsadze, M. Borodovsky, M. Stanke, BRAKER1: unsupervised RNA-Seq-based genome annotation with GeneMark-ET and AUGUSTUS. Bioinformatics 32, 767–769 (2016).

151. K. Hoff, A. Lomsadze, M. Borodovsky, M. Stanke, Whole-genome annotation with BRAKER. 978 Methods Mol Biol (2019).

152. T. Br\uuna, K. J. Hoff, A. Lomsadze, M. Stanke, M. Borodovsky, BRAKER2: automatic eukaryotic genome annotation with GeneMark-EP+ and AUGUSTUS supported by a protein database. NAR Genomics Bioinforma. 3, lqaa108 (2021).

153. A. Lomsadze, P. D. Burns, M. Borodovsky, Integration of mapped RNA-Seq reads into automatic training of eukaryotic gene finding algorithm. Nucleic Acids Res. 42, e119–e119 (2014).

154. S. F. Altschul, W. Gish, W. Miller, E. W. Myers, D. J. Lipman, Basic local alignment search tool. J. Mol. Biol. 215, 403–410 (1990).

155. P. Jones, et al., InterProScan 5: genome-scale protein function classification. Bioinformatics 30, 1236–1240 (2014).

156. D. M. Emms, S. Kelly, OrthoFinder: phylogenetic orthology inference for comparative genomics. Genome Biol. 20, 1–14 (2019).

157. H. Marshall, et al., Parent of origin gene expression in the bumblebee, Bombus terrestris, supports Haig’s kinship theory for the evolution of genomic imprinting. Evol. Lett. 4, 479–490 (2020).

158. B. Langmead, S. L. Salzberg, Fast gapped-read alignment with Bowtie 2. Nat. Methods 9, 357–359 (2012).

159. E. Garrison, G. Marth, Haplotype-based variant detection from short-read sequencing. ArXiv Prepr. ArXiv12073907 (2012).

160. P. Danecek, et al., The variant call format and VCFtools. Bioinformatics 27, 2156–2158 (2011).

161. A. R. Quinlan, I. M. Hall, BEDTools: a flexible suite of utilities for comparing genomic features. Bioinformatics 26, 841–842 (2010).

162. A. Dobin, et al., STAR: ultrafast universal RNA-seq aligner. Bioinformatics 29, 15–21 (2013).

163. F. Krueger, S. R. Andrews, SNPsplit: Allele-specific splitting of alignments between genomes with known SNP genotypes. F1000Research 5 (2016).

164. N. L. Bray, H. Pimentel, P. Melsted, L. Pachter, Near-optimal probabilistic RNA-seq quantification. Nat. Biotechnol. 34, 525–527 (2016).

165. M. D. Robinson, D. J. McCarthy, G. K. Smyth, edgeR: a Bioconductor package for differential expression analysis of digital gene expression data. Bioinformatics 26, 139–140 (2010).

166. K. Riehl, C. Riccio, E. A. Miska, M. Hemberg, TransposonUltimate: software for transposon classification, annotation and detection. Nucleic Acids Res. 50, e64–e64 (2022).

