## Supplementary Materials for "Recent evolution of a maternally-acting sex-determining supergene in a fly with single-sex broods"

This PDF file includes:
Supplementary text 1-7
Supplementary figures 1-4
Supplementary tables 1-4

**Supplementary text 1. *De novo* assembly of reads from X'X individuals.**

Initially, PacBio sequel 3.0 reads from X'X females were assembled *de novo* using wtdbg2 (1) with a lower read length threshold of 2500bp, into 2382 contigs spanning 307Mb with an N50 of 353Kb. The assembly was polished twice with the PacBio reads and three times with 150bp Illumina reads also from X'X individuals using Racon (2). Blobtools v1.1.1 (3) and BUSCO v4 with the insecta_odb10 dataset (4) were used to assess assembly quality and a custom R script was used to remove contaminant and low-coverage contigs. The final BUSCO score was 96.9% (93.5% complete and single-copy BUSCOs, 3.4% complete and duplicated BUSCOs). Scaffolds were assigned to chromosomes based on differences in X0 male and X'X female read coverage. To this end, reads were mapped to the assembly using BWA-MEM (5) and per-based genome coverage was calculated using BEDTools (6). A custom R (7) script was used to assign scaffolds to chromosomes: scaffolds with a relative coverage of 2x in both males and X'X females were assigned as autosomal; scaffolds with a relative coverage of 1x in males and 1-2x in X'X females were assigned to the X chromosome; scaffolds with low (<0.5x) male coverage and 1x coverage in females were assigned to the inversion (**Figure S1**). A custom R script was used to calculate and plot mean coverage across 20Kb windows of the assembly. In total, only 3.63Mb of sequence was assigned as X' inversion sequence using this method. The amount of sequence assigned to the autosomes and X chromosome was similar to that assigned in (8).

**Supplementary text 2. Binning reads with K-mers.**

To assign K-mers as X'-specific, 27-mers were counted in 150bp Illumina read libraries for female-producing female (X'X) and male (X0, from 9) datasets using KMC (10). 27-mers with a frequency of 0 in the X0 library and >5 in the X'X library (to exclude K-mers containing read errors) were assigned as X'-specific; this decision was based on the distribution of 27-mer frequencies visible in **Figure 4.** The cookiecutter (11) option 'extract' was used to pull reads from the raw read file. Cookiecutter creates two separate files: one for all reads that contain a matching K-mer and one containing reads whose mate contained a matching K-mer; both files were combined into one because a read should originate from the same chromosome as its mate. This was done for both 150bp and 75bp Illumina libraries that were available. Extracting putative X' reads with the X'-specific 27-mer library resulted in around 10-12% of reads being pulled; this was roughly in line with the expected size of the X' relative to the rest of the haploid genome (around 40-60Mb in the expected 330-350Mb X'X genome, or 12-17%). When attempting the same process for PacBio sequel 3.0 error-prone reads, significantly more reads were pulled than expected: around 23% of X'-specific reads. We also identified putative autosomal- and X-specific K-mers based on the frequencies visible in **Figure 4**, similar to the methods used in (9), and used these to extract putative autosomal and X reads. For Illumia reads the results were broadly as expected: around 60% for autosomal reads and 13% for X reads. Longer K-mer lengths resulted in very few reads being extracted. It is likely that shorter K-mers occur more often by chance in long reads and result in undesired reads being pulled, while longer K-mers are unlikely to have any matches due to the high error rate of the PacBio reads. For this reason, we decided to competitively map the PacBio reads post-assembly and use them for plugging gaps (**Figure S3**).

**Supplementary text 3. Identification of degraded genes of interest.**

To identify degraded genes that may be of interest, we searched among functional annotations of genes that we classified as degraded using keywords. To identify genes potentially involved in sex determination by chromosome elimination, the keywords used were: 'chromosome', 'chromatid', 'chromatin', 'cohesion', 'condensation', 'segregation', 'centromere', 'centrosome' and 'spindle'. To identify candidates responsible for the wavy wing mutation, we searched for the keyword 'wing'. To identify potential genes with roles in male fertility, the keywords used were 'sex', 'sperm', 'fertility', and 'meiosis'.

**Supplementary text 4. Fly husbandry.**

*Bradysia coprophila* flies have been reared in laboratory conditions since the 1920s (12). Colonies can be successfully maintained in a variety of ways, but are healthiest when kept at around 18-21°C and around 70% relative humidity, and when conditions are not too crowded. At Edinburgh University, colonies are maintained using matings between a single female and two males (to protect against sterility) and are reared on biological agarose in 25mm x 95mm glass vials at 18°C at xxx relative humidity. At the Carnegie Institution for Science in Baltimore, colonies are maintained using mass matings controlled for the sex of offspring (5 males plus 5 female-producing or 5 male-producing females) and are reared in 60z Square Bottom Polypropylene Bottles (Genesee Scientific) capped with Flystuff Flugs (Genesee Scientific) at 21°C and >70% relative humidity. At both institutions, flies are kept on 2.2% Bactoagar and are fed a mixture of yeast, powered mushroom, powdered spinach or nettle and ground straw while larvae.

Hungerford (13) first reported the generation time of *B. coprophila* (from fertilized egg to eclosion) at between 24 and 32 days. Rieffel and Crouse (14) timed developmental stages so as to reflect the minimum length of each stage in ideal conditions, i.e. when crowding, food scarcity, competition and temperature do not interfere with normal development. These minimum developmental stages summed together are 24.6 days for males and 26.6 days for females. Having raised generations of *B. coprophila* in the lab for several years, the authors of this paper have observed the minimum generation time as approximately 28 days, though adults from a single progeny often continue to eclose for up to a week. The generation time fluctuates with temperature: the authors have previously raised *B. coprophila* at 25°C, which speeds up development by at least a few days, and it has previously been noted that rearing *B. coprphila* and *B. ocellaris* at temperatures as low as 12°C is possible but significantly delays development (15, 16). *B. coprophila* have also been observed to survive up to several months when refrigerated (R. B. Baird, C. N. Hodson & L. Ross, unpublished observations; 8). This potential adaptation to cold temperatures may suggest that larvae lie dormant for long periods in winter, which might increase the average generation time. Harsher conditions such as crowding and food scarcity do also appear to extend the generation time beyond 40 days (J. M. Urban & R. B. Baird, unpublished observations). On the other hand, the generation time may be shorter in regions with warmer climates. Observations of other, closely-related species report varying generation times: 26-48 days or *B. odoriphaga* (17), 14 days for *B. impatiens* (18), 25 days at 20-25°C or 3-4 weeks at 22-24°C for *B. paupera* (19) and 26-28 days at 25°C for *B. difformis* (20); see (21) for a review). Overall, it is difficult to estimate the generation time that best approximates *B. coprophila* development in nature, but in light of the literature cited here and observations made by the authors we decided to use a generation time 24-40 days to calculate divergence between the X and X' chromosomes in *B. coprophila* in years.

**Supplementary text 5. Genome annotation.**

The RNAseq data used in annotation of the genome (including autosomes II, III, IV, X and the X′ sequence) was obtained from publicly available data previously produced by Urban *et al.* (8, 22, 23) and unpublished datasets produced by the Ross Lab. This included 18 RNAseq datasets from four life stages (3 x male and 3 x female embryos, 2 x male and 2 x female larval, 2 x male and 2 x female pupal, 2 x male and 2 x female adult, (8), 15 datasets from dissected larval salivary glands (pooled individuals, (22), 6 datasets from radiated larvae (pooled female individuals, (23), 6 datasets from male and female somatic tissue (3 x male, 3 x female larval/early pupal), as well as 12 datasets from early (0-8h) embryos (6 x male, 6 x female, unpublished data). All RNAseq datasets were aligned to the genome using STAR (24) prior to being fed to BRAKER2. Homology-based datasets included all OrthoDB v10 Diptera protein sequences (751660 sequences,(25), all Uniprot Diptera protein sequences (7053 sequences, (26) and all Refseq Diptera protein sequences including all non-canonical isoforms (1575334 sequences, (27).

Functional information for the 26887 protein sequences in the BRAKER2 gene annotation set was obtained by finding the best BLASTP (28) hits in several protein databases and with InterProScan. Specifically, all BLASTP hits (-word_size 3 -evalue 1e-2) for all 26887 proteins were found in (i) the *Drosophila melanogaster* r6.45 (29) proteome, (ii) Non-redundant UniProtKB/SwissProt (30) protein sequences (ftp://ftp.ncbi.nlm.nih.gov/blast/db/swissprot.tar.gz), (iii) the NCBI Landmark database for SmartBLAST (ftp://ftp.ncbi.nlm.nih.gov/blast/db/landmark.tar.gz), (iv) the NCBI RefSeq (31) protein database (ftp://ftp.ncbi.nlm.nih.gov/blast/db/refseq_protein.*.tar.gz), (v) the NCBI Non-Redundant (nr) protein database (ftp://ftp.ncbi.nlm.nih.gov/blast/db/nr.*.tar.gz), (vi) OrthoDB v10 (25), (vii) and gene annotations from the original *Bradysia coprophila* reference genome (40). The best BLASTP hit for all 26887 proteins in each database was found by taking the hit with the highest bitscore. Hits with bitscores lower than 50 or e-values higher than 0.0005 were removed from consideration. The 26887 protein sequences were also extensively analyzed using the InterPro (32) protein family and domain database and InterProScan version 5.56-89.0 (-dp -iprlookup -goterms -f tsv,xml,json,gff3, (33) allowing all analyses to be run: CDD, Coils, Gene3D, Hamap, MobiDBLite, PANTHER, Pfam, Phobius, PIRSF, PIRSR, PRINTS, ProSitePatterns, ProSiteProfiles, SFLD, SignalP, SMART, SUPERFAMILY, TIGRFAM, TMHMM. Software licenses were obtained where relevant (SignalP, Phobius, TMHMM).

**Supplementary text 7. Transposable element annotation.**

To annotate transposable elements (TEs) in the genome, the module reasonaTE of the TranposonUltimate v1.03 pipeline (34) was used (**S7 Text**). Combining the different annotation approaches and sensitivities of RepeatMasker (v4.1.2, (35), RepeatModeler (v2.0.2a, (36), LTRharvest (2.9.0, (37), TIRvish (v.1.10.5, (38), SINE-Scan (v1.1.2, (39), HelitronScanner (v1.0.0, (40), MiteFinderII (v1.0.0, (41), MITE-Tracker (v1.0.1, (42), TransposonPSI (v1.0.0, https://transposonpsi.sourceforge.net/) and NCBICDD1000 (v1.0.1, (43) it enables in-depth analysis of various classes of TEs in the genome. R Studio (7) was used to plot TE distribution across chromosomes.

**Supplementary text 6. Data used to examine dosage compensation.**

To investigate potential dosage compensation (DC) of X-linked genes with degraded X′ homologs, we used RNAseq data from early (0-8h) embryos. This dataset was generated by R. B. Baird for the purpose of another, unpublished study. Adult females were mated with males (generally ~10 females with ~3 males in a tube), and after ~18-24 hours, females were pinned to a petri dish containing 2.2% Bactoagar and egg laying was induced by crushing the head slightly. An entire clutch (50-100 eggs) is usually laid within the next 2 hours. Three replicates from two time stages were collected: 0-4h and 4-8h after egg deposition (AED), for the eggs of each female genotype (X′X and XX), resulting in 12 samples in total. For each sample, around ~600-800 eggs pooled from ~15 individuals were collected and sequenced for 15Gb (50m reads) of paired-end 150bp RNAseq reads on the Illumina Novaseq S1 platform.

We used these data to examine expression of X-linked genes in X′X versus XX females because data for female adults (or other developmental stages) separated by genotype are not yet available, and these early embryos are likely to contain exclusively maternal transcripts. In *Drosophila*, zygotic genome activation (ZGA) occurs in two waves at mitotic cycle 8 (60 genes) and following cellularization at mitotic cycle 14 (over 1000 genes, 22). De Saint Phalle & Sullivan (45) report that cellularization in *B. coprophila* occurs during interphase of nuclear cycle 11, which begins at 9.3 +/- 1.1 hours AED. Cell cycle 12 begins at 11.4 +/- 1.4 hours AED. However, as in *Drosophila*, de Saint Phalle & Sullivan (45) report that the germline nuclei cellularize slightly earlier, at cell cycle 7, which occurs around 4.2 +/- 0.5 hours AED. Nonetheless, if we assume that, as in *Drosophila*, ZGA occurs approximately when cellularization occurs, ZGA should begin for the majority of somatic genes by 12 hours AED.

Upon splitting RNAseq reads from the eggs of X'X mothers into X´ and X reads (see methods in main text), similar proportions of reads were assigned to the X´ (5.67%) and X (6.17%) aside from one outlier sample (eggs of X´X females, 4-8h, replicate 2), which had 7.97% of reads assigned to the X and 3.61% assigned to the X´). This sample was excluded from further analysis. If transcripts were zygotic rather than maternal, then the number of X reads assigned should be three-fold higher for the X, because in the pooled eggs, maternal transcripts from X´X mothers will originate from the X´ and X in equal proportions. However, following ZGA, three times as many X transcripts should be produced for every X´ transcript, owing to half the eggs being XX and the other half X´X.

**Supplementary table 1.** Assembly statistics and chromosome anchorage for the *B. coprophila* genome assembled *de novo* from PacBio long reads from X'X individuals.

| Partition | Length (Mb) | N scaffolds | N50 (Kb) | Largest scaffold (Kb) |
|---|---|---|---|---|
| Autosomes | 215.15 | 1123 | 423.42 | 2141.89 |
| X chromosome | 72.84 | 864 | 205.13 | 995.08 |
| Inversion | 3.63 | 259 | 17.69 | 73.99 |
| Total | 291.63 | 2246 | 342.36 | |

**Supplementary table 2.** Number of genes within the X' supergene sequence with each type of predicted pseudogenizing mutation.

| Mutation type | N genes |
|---|---|
| Stop codon gained | 37 |
| Stop codon lost | 10 |
| Start codon lost | 10 |
| Frameshift | 113 |
| Frameshift and stop codon gained | 15 |
| Frameshift and stop codon lost | 6 |
| Frameshift and start codon lost | 8 |
| Frameshift and stop codon gained and stop codon lost | 2 |
| Frameshift and stop codon gained and start codon lost | 1 |

**Supplementary table 3.** Functions of pseudogenized genes of interest.

| Gene ID | Functionality | Annotation(s) |
|---|---|---|
| jg4815 | nonfunctional_disrupted | Similar to how Protein held out wings |
| jg6281 | nonfunctional_silenced | Similar to su(Hw) Protein suppressor of hairy wing |
| jg5727 | nonfunctional_disrupted | Similar to su(Hw) Protein suppressor of hairy wing |
| jg7913 | nonfunctional_disrupted | Similar to su(Hw) Protein suppressor of hairy wing |
| jg7400 | nonfunctional_disrupted | Similar to stan Protocadherin-like wing polarity protein |
| jg6904 | nonfunctional_disrupted | Similar to Etl1 SWI/SNF-related matrix-associated actin-dependent regulator of chromatin subfamily A containing DEAD/H box 1 homolog; Similar to Marcal1 SWI/SNF-related matrix-associated actin-dependent regulator of chromatin subfamily A-like protein 1 |
| jg6344 | nonfunctional_disrupted | Similar to Etl1 SWI/SNF-related matrix-associated actin-dependent regulator of chromatin subfamily A containing DEAD/H box 1 homolog; Similar to Marcal1 SWI/SNF-related matrix-associated actin-dependent regulator of chromatin subfamily A-like protein 1 |
| jg7872 | nonfunctional_silenced | Similar to dsx Protein doublesex |
| jg6244 | nonfunctional_disrupted | Similar to Crocc Rootletin; Ciliary rootlet component, centrosome cohesion |
| jg6043 | nonfunctional_silenced | Similar to TRAF3IP1 TRAF3-interacting protein 1; Microtubule-binding protein MIP-T3 C-terminal region |
| jg5804 | nonfunctional_disrupted | Similar to ncd Protein claret segregational |

**Supplementary table 4.** Upregulated X-linked genes with single-copy X'-linked homologs in X'X females.

| Gene ID | Annotation(s) |
|---|---|
| jg4857 | Similar to PARP3 Protein mono-ADP-ribosyltransferase PARP3 |
| Jg5988 | Similar to srfbp1 Serum response factor-binding protein 1;<br>Similar to F52C9.6 Putative uncharacterized transposon-derived protein F52C9.6 |
| Jg6324 | Protein of unknown function |
| Jg7315 | Similar to eff Ubiquitin-conjugating enzyme E2-17 kDa |

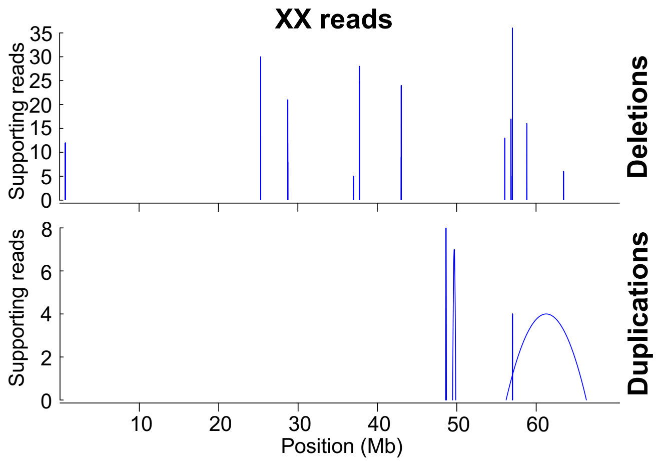


**Supplementary figure 1.** Structural variant calls from the XX genotype from Illumina 75bp paired-end short reads. These calls serve as an extra control along with calls from the X0 genotype, which, when constrasted with calls from the X'X genotype, provide further support that the X' is enriched for complex rearrangements.

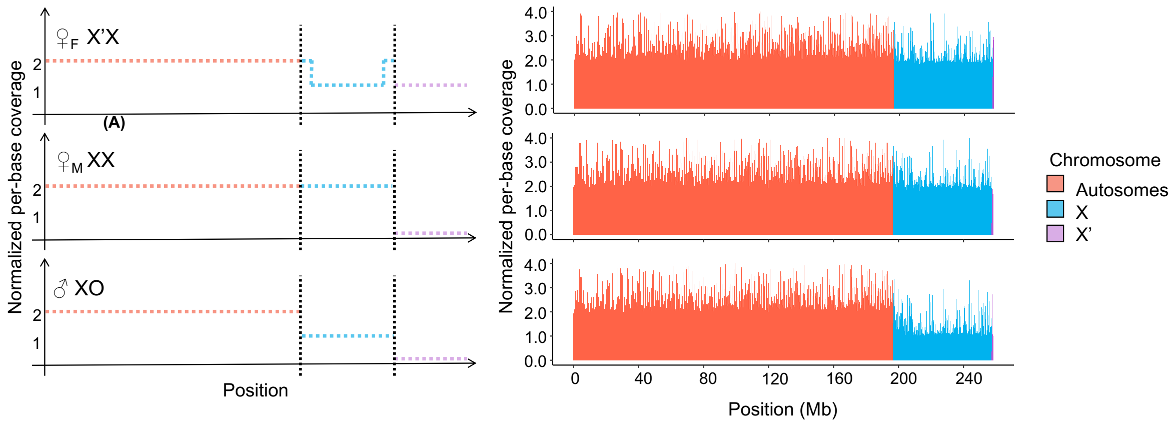


**Supplementary figure 2.** Expected (A) versus observed (B) differences in genomic coverage across the genome assembled *de novo* from PacBio reads from X'X individuals, indicating that the vast majority of reads from the X and X' chromosomes collapsed together upon assembly. Note that the assembly is 291.63 Mb in length but appears shorter in the figure above because some scaffolds were shorter than the 20 Kb windows across which coverage was calculated, so such scaffolds were not included.

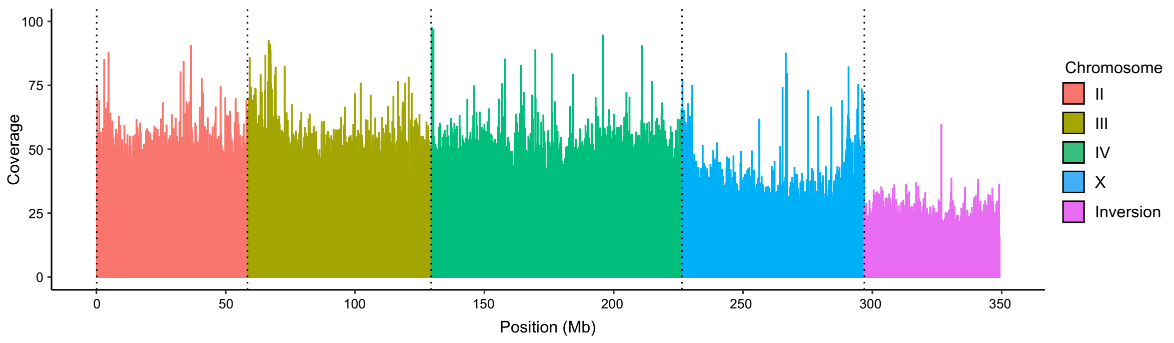


**Supplementary figure 3.** PacBio long reads from X'X individuals mapped to the X'X assembly. Now that the X and X' inversion are separately assembled, PacBio reads map to the correct chromosomes with approximately expected coverage and could thus be used to fill some remaining gaps in the assembly of the X' inversion.

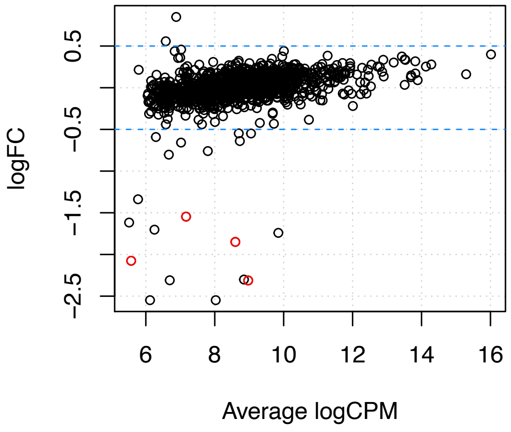


**Supplementary figure 4.** Differential expression smear plot for X-linked genes with single-copy X' homologs between the two types of females (XX versus X'X). Negative fold change (FC) represents upregulation of the gene copy in X'X females; positive FC represents upregulation of the gene in XX females. Transcript counts have been normalized such that overall X in X'X females expression equals that of XX females. Genes with significant fold change and significant adjusted P values are coloured red.

### Supplementary references


1. J. Ruan, H. Li, Fast and accurate long-read assembly with wtdbg2. *Nat. Methods* **17**, 155–158 (2020).

2. R. Vaser, I. Sović, N. Nagarajan, M. Šikić, Fast and accurate de novo genome assembly from long uncorrected reads. *Genome Res.* **27**, 737–746 (2017).

3. D. R. Laetsch, M. L. Blaxter, BlobTools: Interrogation of genome assemblies. *F1000Research* **6**, 1287 (2017).

4. M. Seppey, M. Manni, E. M. Zdobnov, "BUSCO: Assessing Genome Assembly and Annotation Completeness" in *Gene Prediction: Methods and Protocols*, M. Kollmar, Ed. (Springer New York, 2019), pp. 227–245.

5. H. Li, Aligning sequence reads, clone sequences and assembly contigs with BWA-MEM. *ArXiv Prepr. ArXiv13033997* (2013).

6. A. R. Quinlan, I. M. Hall, BEDTools: a flexible suite of utilities for comparing genomic features. *Bioinformatics* **26**, 841–842 (2010).

7. R Core Team, *R: A Language and Environment for Statistical Computing* (R Foundation for Statistical Computing, 2022).

8. J. M. Urban, *et al.*, High contiguity de novo genome assembly and DNA modification analyses for the fungus fly, *Sciara coprophila*, using single-molecule sequencing. *BMC Genomics* **22**, 643 (2021).

9. C. N. Hodson, K. S. Jaron, S. Gerbi, L. Ross, Gene-rich germline-restricted chromosomes in black-winged fungus gnats evolved through hybridization. *PLOS Biol.* **20**, e3001559 (2022).

10. M. Kokot, M. Długosz, S. Deorowicz, KMC 3: counting and manipulating k-mer statistics. *Bioinformatics* **33**, 2759–2761 (2017).

11. E. Starostina, G. Tamazian, P. Dobrynin, S. O'Brien, A. Komissarov, "Cookiecutter: a tool for kmer-based read filtering and extraction" *Bioinformatics* (2015).

12. Chas. W. Metz, Chromosomes and Sex in *Sciara*. *Science* **61**, 212–214 (1925).

13. H. Hungerford, Sciara maggots injurious to potted plants. *J. Econ. Entomol.* **9**, 538–549 (1916).

14. S. Rieffel M., H. V. Crouse, The elimination and differentiation of chromosomes in the germ line of *Sciara*. *Chromosoma* **19**, 231–276 (1966).

15. H. Smith-Stocking, Genetic studies on selective segregation of chromosomes in *Sciara coprophila* Lintner. *Genetics* **21**, 421–443 (1936).

16. R. G. Nigro, M. C. C. Campos, A. L. P. Perondini, Temperature and the progeny sex-ratio in *Sciara ocellaris* (Diptera, Sciaridae). *Genet. Mol. Biol.* **30**, 152–158 (2007).

17. W. Li, *et al.*, Effects of Temperature on the Age-Stage, Two-Sex Life Table of *Bradysia odoriphaga* (Diptera: Sciaridae). *J. Econ. Entomol.* **108**, 126–134 (2015).

18. M. K. Kennedy, A culture method for *Bradysia impatiens* (Diptera: Sciaridae). *Ann. Entomol. Soc. Am.* **66**, 1163–1164 (1973).

19. J. Mansilla, M. Pastoriza, R. Pérez, Study on biology and control of *Bradysia paupera* Tuomikoski (= *Bradysia difformis* Frey)(Diptera: Sciaridae). *Bol. Sanid. Veg. Plagas Esp.* (2001).

20. E. Villanueva-Sánchez, S. Ibáñez-Bernal, J. Lomelí-Flores, J. Valdez-Carrasco, Identificación y caracterización de la mosca en el cultivo de nochebuena (*Euphorbia pulcherrima*) en el centro de México. *Acta Zool. Mex.* **29**, 363–375 (2013).

21. A. Katumanyane, T. Ferreira, A. P. Malan, A Review of *Bradysia* spp. (Diptera: Sciaridae) as Pests in Nursery and Glasshouse Crops, With Special Reference to Biological Control Using Entomopathogenic Nematodes. *Afr. Entomol.* **26**, 1–13 (2018).

22. J. Urban, *et al.*, Re-replication Origins in Sciara DNA Puffs Revealed by New and Old Genomic Technologies including Nanopore Sequencing. *FASEB J.* **29**, 561–9 (2015).

23. J. M. Urban, *et al.*, "*Sciara coprophila* larvae upregulate DNA repair pathways and downregulate developmental regulators in response to ionizing radiation" *Genomics* (2021).

24. A. Dobin, *et al.*, STAR: ultrafast universal RNA-seq aligner. *Bioinformatics* **29**, 15–21 (2013).

25. E. V. Kriventseva, *et al.*, OrthoDB v10: sampling the diversity of animal, plant, fungal, protist, bacterial and viral genomes for evolutionary and functional annotations of orthologs. *Nucleic Acids Res.* **47**, D807–D811 (2019).

26. U. Consortium, UniProt: a hub for protein information. *Nucleic Acids Res.* **43**, D204–D212 (2015).

27. K. D. Pruitt, T. Tatusova, D. R. Maglott, NCBI reference sequences (RefSeq): a curated non-redundant sequence database of genomes, transcripts and proteins. *Nucleic Acids Res.* **35**, D61–D65 (2007).

28. S. F. Altschul, W. Gish, W. Miller, E. W. Myers, D. J. Lipman, Basic local alignment search tool. *J. Mol. Biol.* **215**, 403–410 (1990).

29. G. dos Santos, *et al.*, FlyBase: introduction of the *Drosophila melanogaster* Release 6 reference genome assembly and large-scale migration of genome annotations. *Nucleic Acids Res.* **43**, D690–D697 (2014).

30. Uniprot Consortium, UniProt: the universal protein knowledgebase in 2021. *Nucleic Acids Res.* **49**, D480–D489 (2020).

31. N. A. O'Leary, *et al.*, Reference sequence (RefSeq) database at NCBI: current status, taxonomic expansion, and functional annotation. *Nucleic Acids Res.* **44**, D733–D745 (2015).

32. M. Blum, *et al.*, The InterPro protein families and domains database: 20 years on. *Nucleic Acids Res.* **49**, D344–D354 (2020).

33. P. Jones, *et al.*, InterProScan 5: genome-scale protein function classification. *Bioinformatics* **30**, 1236–1240 (2014).

34. K. Riehl, C. Riccio, E. A. Miska, M. Hemberg, TransposonUltimate: software for transposon classification, annotation and detection. *Nucleic Acids Res.* **50**, e64–e64 (2022).

35. A. Smit, R. Hubley, P. Green, RepeatMasker Open-4.0. 2013–2015 (2015).

36. J. M. Flynn, *et al.*, RepeatModeler2 for automated genomic discovery of transposable element families. *Proc. Natl. Acad. Sci.* **117**, 9451–9457 (2020).

37. D. Ellinghaus, S. Kurtz, U. Willhoeft, LTRharvest, an efficient and flexible software for de novo detection of LTR retrotransposons. *BMC Bioinformatics* **9**, 1–14 (2008).

38. G. Gremme, S. Steinbiss, S. Kurtz, GenomeTools: a comprehensive software library for efficient processing of structured genome annotations. *IEEE/ACM Trans. Comput. Biol. Bioinform.* **10**, 645–656 (2013).

39. H. Mao, H. Wang, SINE_scan: an efficient tool to discover short interspersed nuclear elements (SINEs) in large-scale genomic datasets. *Bioinformatics* **33**, 743–745 (2017).

40. W. Xiong, L. He, J. Lai, H. K. Dooner, C. Du, HelitronScanner uncovers a large overlooked cache of Helitron transposons in many plant genomes. *Proc. Natl. Acad. Sci.* **111**, 10263–10268 (2014).

41. J. Hu, Y. Zheng, X. Shang, MiteFinderII: a novel tool to identify miniature inverted-repeat transposable elements hidden in eukaryotic genomes. *BMC Med. Genomics* **11**, 51–59 (2018).

42. J. M. Crescente, D. Zavallo, M. Helguera, L. S. Vanzetti, MITE Tracker: an accurate approach to identify miniature inverted-repeat transposable elements in large genomes. *BMC Bioinformatics* **19**, 1–10 (2018).

43. S. Lu, *et al.*, CDD/SPARCLE: the conserved domain database in 2020. *Nucleic Acids Res.* **48**, D265–D268 (2020).

44. E. Darbo, C. Herrmann, T. Lecuit, D. Thieffry, J. van Helden, Transcriptional and epigenetic signatures of zygotic genome activation during early drosophila embryogenesis. *BMC Genomics* **14**, 226 (2013).

45. B. de Saint Phalle, W. Sullivan, Incomplete sister chromatid separation is the mechanism of programmed chromosome elimination during early *Sciara coprophila* embryogenesis. *Development* **122**, 3775–3784 (1996).
